# Neurally-constrained modeling of human gaze strategies in a change blindness task

**DOI:** 10.1101/663989

**Authors:** Akshay Jagatap, Hritik Jain, Simran Purokayastha, Devarajan Sridharan

**Affiliations:** Centre for Neuroscience, Indian Institute of Science, Bangalore 560012, India; Department of Psychology, New York University, New York 10003, USA; Computer Science and Automation, Indian Institute of Science, Bangalore 560012, India

**Author notes:** Corresponding author: Devarajan Sridharan, Centre for Neuroscience, Rm F104 Indian Institute of Science, C. V. Raman Avenue, Bangalore 560012, India.

**Keywords:** Attention, change detection, visual search, saccades, Bayesian behavior models, SPRT

## Abstract

Visual attention enables us to engage selectively with the most important events in the world around us. Yet, sometimes, we fail to notice salient events. “Change blindness” – the surprising inability to detect and identify salient changes that occur in flashing visual images – enables measuring such failures in a laboratory setting. We discovered that human participants (n=39) varied widely (by two-fold) in their ability to detect changes when tested on a laboratory change blindness task. To understand the reasons for these differences in change detection abilities, we characterized eye-movement patterns and gaze strategies as participants scanned these images. Surprisingly, we found no systematic differences between scan paths, fixation maps or saccade patterns between participants who were successful at detecting changes, versus those who were not. Yet, two low-level gaze metrics – the mean fixation duration and the variance of saccade amplitudes – systematically predicted change detection success. To explain the mechanism by which these gaze metrics could influence performance, we developed a neurally constrained model, based on the Bayesian framework of sequential probability ratio testing (SPRT), which simulated gaze strategies of successful and unsuccessful observers. The model’s ability to detect changes varied systematically with mean fixation duration and saccade amplitude variance, closely mimicking observations in the human data. Moreover, the model’s success rates correlated robustly with human observers’ success rates, across images. Our model explains putative human attention mechanisms during change blindness tasks and provides key insights into effective strategies for shifting gaze and attention for artificial agents navigating dynamic, crowded environments.

**Author Summary:** Our brain has the remarkable capacity to pay attention, selectively, to the most important events in the world around us. Yet, sometimes, we fail spectacularly to notice even the most salient events. We tested this phenomenon in the laboratory with a change-blindness experiment, by having participants freely scan and detect changes across discontinuous image pairs. Participants varied widely in their ability to detect these changes. Surprisingly, their success correlated with differences in low-level gaze metrics. A Bayesian model of eye movements, which incorporated neural constraints on stimulus encoding, could explain the reason for these differences, and closely mimicked human performance in this change blindness task. The model’s gaze strategies provide relevant insights for artificial, neuromorphic agents navigating dynamic, crowded environments.

## Introduction

We live in a rapidly-changing world. For adaptive survival, our brains must possess the ability to identify changing, relevant aspects of our environment and distinguish them from static, irrelevant ones. For example, when driving down a busy road, it is essential to identify changing aspects of the visual scene that are relevant for safe navigation, such as tracking movements of a vehicle shifting lanes in front or looking out for pedestrian traffic, while ignoring irrelevant aspects, such as objects by the wayside. Visual attention is the essential cognitive capacity that enables us to efficiently identify and respond to only the most relevant events, at each moment in time, to guide behavior (Carrasco 2011).

Yet, our capacity for attention possesses key limitations. One such limitation is revealed by the phenomenon of “change blindness”, in which observers fail to detect obvious changes in a sequence of visual images with intervening discontinuities (Simons and Ambinder 2005; Simons and Rensink 2005). It has been suggested that observers’ lapses with detecting changes are not due to impaired visual processing of the changing aspects of the image: changes are detected readily enough if the images occur in succession, with no intervening discontinuities. Rather, these lapses occur if the changes fail to attract attention, for example if the image is blanked before the change or in the presence of concurrent distractors that render it challenging to detect and localize the change. Change blindness, therefore, provides a useful framework for studying visual attention mechanisms and its lapses (Rensink, O’Regan, and Clark 1997), with important real-world implications. In fact, observers’ ability to detect changes in change blindness tasks has been linked to their driving experience (Crundall 2009; Zhao et al. 2014) and ability to drive safely (Beanland, Filtness, and Jeans 2017).

In the laboratory, change blindness is tested with tasks of the type shown in Figure 1A. A pair of images, typically crowded scenes that differ in one important detail, are presented alternately, with an intervening blank (“flicker” paradigm) (Simons and Ambinder 2005; Simons and Rensink 2005). Participants scan the images, by moving their eyes to different regions of the image, in sequence, to locate and identify the changing object or feature. Because shifts of attention are concomitant with eye movements across a visual scene (Deubel and Schneider 1996; Hoffman and Subramaniam 1995), an individual’s capacity for selective attention is likely closely linked with her/his ability to perform well in such change blindness tasks. Consistent with this hypothesis, previous studies (Rensink, O’Regan, and Clark 1997) have shown that directing attention toward the location of change – even covertly, i.e., without moving the eyes – renders the change more easily detectable.

**Figure 1.**
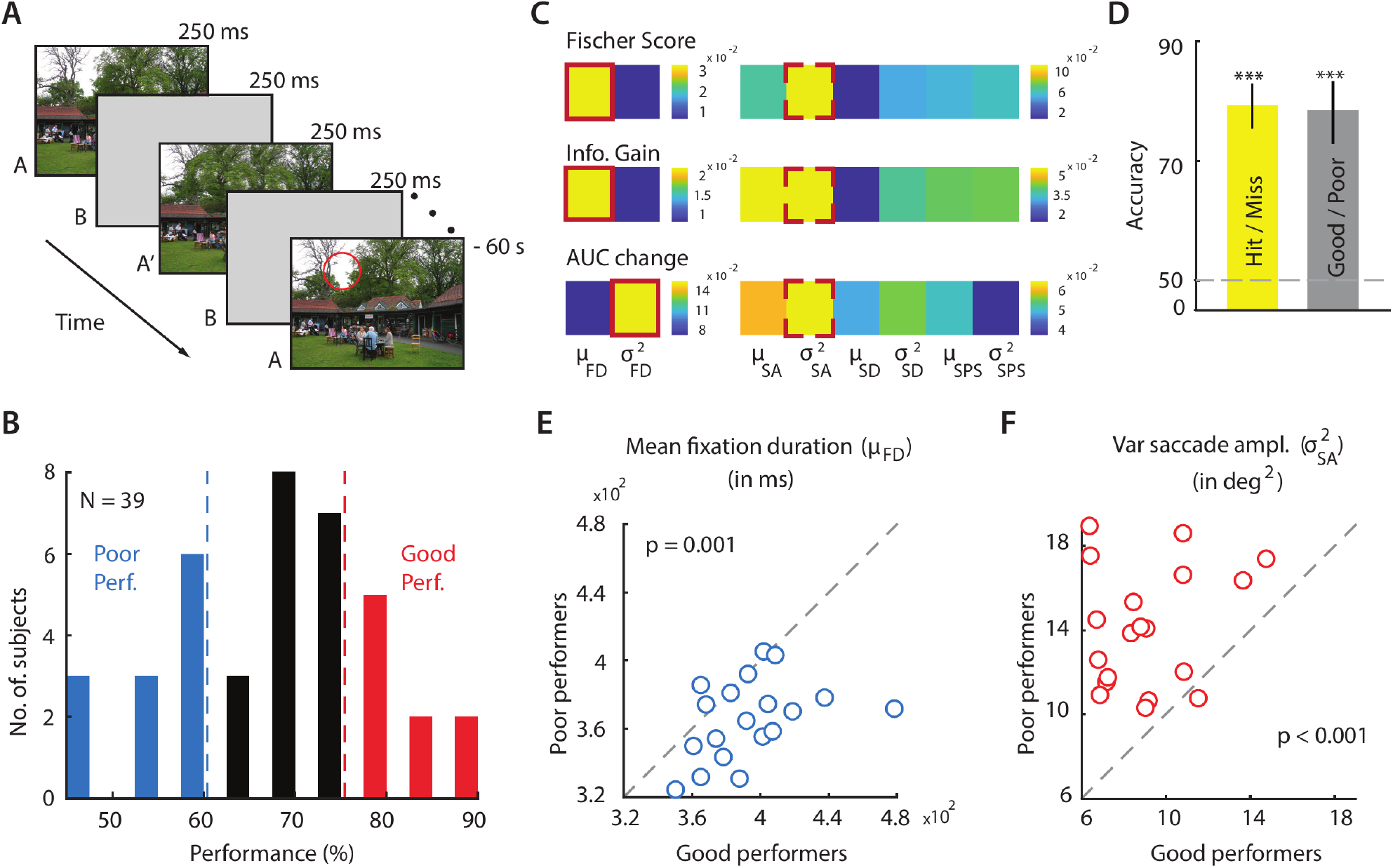
Gaze metrics predictive of success in a change blindness task. **A.** Schematic of a change blindness experiment trial, comprising a sequence of alternating images (A, A’), displayed for 250 ms each, with intervening blank frames (B) also displayed for 250 ms (“flicker” paradigm), repeated for 60 s. Red circle: Location of change (not actually shown in the experiment). All 20 change image pairs tested are shown in SI Figure S1. **B.** Distribution of success rates of n=39 participants in the change blindness experiment. Red and blue bars: good performers (top 30^th^ percentile; n=10) and poor performers (bottom 30th percentile; n=12), respectively. Dashed vertical lines: Cut-off values of success rates for classifying good versus poor performers. **C.** Feature selection measures for identify the most informative features that distinguish good from poor performers. Top row: Fisher score; middle row: Information gain; bottom row: Change in area-under-the-curve (AUC) (see text for details). Brighter colors indicate more informative features. Solid red outline: most informative feature in the fixation feature subgroup (left); dashed red outline: most informative feature in the saccade feature subgroup (right). FD - fixation duration; SD - saccade duration; SA - saccade amplitude, SPS - saccade peak speed. μ and σ^2^ denote mean and variance of the respective parameter. **D.** Classification accuracy, quantified with area-under-the-curve (AUC), for classifying trials as hits versus misses (left bar) and performers as good versus poor (right bar), obtained with a support vector machine classifier. Dashed horizontal line: Classification accuracy at chance (50%). p-value corresponds to significance level of classification accuracy above chance (permutation test; ***p < 0.001). Error bars: Clopper-Pearson binomial confidence intervals. **E.** Distribution of mean fixation duration (μ_FD_, in milliseconds) across 19 change images for good performers (x-axis) versus poor performers (y-axis); one change image pair, successfully detected by all performers, was not included in these analyses (see text). Each data point denotes average value of μ_FD_, across each category of performers, for each image tested. Dashed diagonal line: line of equality. p-value corresponds to significant difference in mean fixation duration between good and poor performers **F.** Same as in E, but comparing variance of saccade amplitudes (in squared degrees of visual angle) across good versus poor performers. Other conventions are the same as in panel E.

While many previous studies have investigated the phenomenon of change blindness itself (Droll, Gigone, and Hayhoe 2007; O’regan, Rensink, and Clark 1999; Tse 2004), very few studies have directly identified gaze related factors that determine observers’ success in change blindness tasks (Rensink, O’Regan, and Clark 1997). In this study, we tested 44 participants in a change blindness experiment comprising a sequence of 20 image pairs (Fig. 1A). Surprisingly, participants differed widely (by over 2-fold) in their success with detecting changes. To understand the mechanisms underlying these striking differences in performance, we collected high-resolution eye-tracking data, and analyzed these data with machine learning approaches to identify key eye-movement patterns and gaze metrics that were predictive of change detection success. We discovered that two key gaze metrics -- mean fixation duration and the variance in the amplitude of saccades -- were consistently predictive of participants’ success.

To understand the predictive power of these gaze metrics, we developed a model of eye-movements based on the Bayesian framework of sequential probability ratio test (SPRT) (Bogacz et al. 2006; Gold and Shadlen 2000; Smith and Ratcliff 2004; Wald 1973) that also incorporated constraints on neural stimulus encoding (e.g. bounded firing rates, Poisson variance, foveal magnification, saliency computation; Gold and Shadlen 2007; Kira, Yang, and Shadlen 2015). Despite its simple framework, the model mimicked key aspects of human performance in the change blindness task. The model’s success rates were strongly correlated with human success rates across the images tested. Remarkably, as an emergent feature, the model exhibited systematic differences in change detection success rates when fixation duration and saccade amplitude parameters were varied, in a manner closely resembling human data. Our study identifies putative mechanisms for successful gaze strategies in change blindness tasks, which may also provide key insights for designing algorithms for scanning and deploying attention, for autonomous agents navigating crowded environments.

## Results

### Low-level gaze metrics distinguish good from poor performers

39 participants performed a change blindness task (Fig. 1A). Each experimental session consisted of a sequence of trials with a different pair of images tested on each trial. Images tested included cluttered, indoor or outdoor scenes (SI Fig. S1; SI Table S1, SI Videos; Methods). Each trial began when subjects fixated continually on a central cross for 3 seconds; this was done to ensure uniformity of gaze origin across subjects, before they began their search for the change. This was followed by the presentation of the change blindness image pair: alternating frames of two images, separated by intervening blank frames (250 ms each, Fig. 1A). Of the image pairs tested, 20 were “change” image pairs, which differed from each other in one of three key aspects (SI Fig. S1): i) change in the size of an object; ii) change in the color of an object or iii) change involving the appearance (or disappearance) of an object (SI Fig. S1). The remaining (either 6 or 7 pairs across subjects) were “catch” image pairs, which comprised of identical images without any difference between them; data from these “catch” trials were not analyzed for this study (Methods). Change and catch image pairs were interleaved and tested in the same pseudorandom order across subjects. Subjects were permitted to freely scan the images to detect the change, for up to a maximum of 60 seconds per image pair. They indicated having detected the change by foveating at the location of change for at least 3 seconds. A response was marked as a “hit” if the subject was able to successfully locate the change within 60 seconds, and was marked as a “miss” otherwise.

We observed wide variations across participants, in their ability to detect changes: success rates varied over two-fold – from 45% to 90% – across participants (Fig. 1B). Note that these differences may have occurred for a variety of reasons, including each participant’s level of engagement with the task, their motivation or level of alertness on the testing day and the like, and may not reflect individual traits or capacities for change detection (see Discussion). Nevertheless, we sought to understand if these differences in success rates with detecting changes could be explained by individual differences in gaze strategies when scanning the images.

To answer this question, we first, ranked subjects in order of their success rates. Subjects in the top 30th (n=9) and bottom 30th (n=12) percentiles were labelled as “good” and “poor” performers, respectively (Fig. 1B). For these subjects, we computed four gaze metrics from the eye-tracking data: saccade amplitude, fixation duration, saccade duration and saccade peak speed (see Methods for details). We then computed the mean and variance of these four parameters for each subject and for each trial (image pair). These eight measures were employed as features in a classifier based on support vector machines (SVM) to classify good from poor performers. Because all participants correctly detected the change on one of the image pairs (SI Fig. S1 Image #20), we excluded data from this image pair for these analyses.

Classification accuracy, as quantified by area-under-the-curve (AUC), for distinguishing good from poor performers was 79.9% and significantly above chance (Fig. 1D, p<0.001, permutation test, Methods). We repeated these same analyses by seeking to classify each trial as a hit or miss based on these same eight gaze metric parameters. Classification accuracy was 77.7% and, again, significantly above chance (Fig. 1D, p<0.001). Taken together, these results indicate that fixation- and saccade- related gaze metrics contained sufficient information to accurately classify successful performers in this change blindness experiment.

Next, we sought to identify which gaze metrics were the most informative for classifying good versus poor performers. For this, we computed, first, the pair-wise correlation (across trials) between all eight gaze metrics to identify those that were likely to contribute independently to classification performance. We discovered that the mean and variance of fixation duration were mutually (positively) correlated, but uncorrelated (or anti-correlated) with all other parameters (p<0.01; Bonferroni corrected for multiple comparisons; SI Fig. S2A). On the other hand, mean and variance of saccade amplitude, saccade duration and saccade peak speed were almost all mutually correlated with each other (SI Fig. S2A). We term the former, the fixation feature sub-group, and the latter, the saccade feature subgroup, respectively. Next, we performed feature selection analysis among these two sub-groups of features to identify which metrics, within each sub-group, were the most informative for classifying good versus poor performers. For this, we computed three measures for each feature – Fisher score (Boser, Guyon, and Vapnik 1992), sensitivity analysis (Nasrabadi 2007) and information gain (Yang 1997). A higher value of each feature selection measure reflects a greater importance of the corresponding feature, for classifying good versus poor performers. In the fixation feature sub-group, mean fixation duration was flagged as the most important metric compared to variance of fixation duration, based on two out of the three feature selection measures (Fig. 1C, solid red outline). On the other hand, in the saccade feature sub-group, variance of saccade amplitudes was the single most important metric, based on all three feature selection measures (Fig. 1C, dashed red outline). Confirming the results of feature selection, we found systematic differences in the values of these metrics across classes: while mean fixation duration was significantly higher for good performers, across images (Fig. 1E; p=0.0015, Wilcoxon signed rank test), variance of saccade amplitude was significantly higher for poor performers (Fig. 1F; p<0.001, Wilcoxon signed rank test).

### Good and poor performers exhibit similar fixation and scan patterns

In addition to these fixation and saccade metrics, we compared more complex aspects of eye movements between good and poor performers. Specifically, we asked if scan paths, fixation maps or low-level fixated features differed systematically between good and poor performers.

Scan path data is challenging to compare across individuals because scan paths can vary both in number and nature of image locations sampled, and in the temporal sequence of this sampling. To simplify comparing scan paths across participants, we adopted the following approach: fixation points in each image were clustered across subjects, and scan paths were encoded as strings, for each image and subject, based on the sequence of clusters visited at successive fixations (Methods).

First, we computed the edit distance between scan paths (Cristino et al. 2010), for each image across every pair of participants, and tested if the median of these distributions was significantly different between good and poor performers (Methods). Briefly, the edit distance provides an intuitive measure of the dissimilarity between two strings. It corresponds to the minimal number of “edit” operations -- insertions, deletions or substitutions -- that are necessary to transform one string into the other. No significant differences were found between the medians of edit distance distributions of scan paths across good and poor performers (Fig. 2D, p=0.14, Wilcoxon signed rank test). To further confirm the lack of differences, we compared the median intra-category edit distance (among the good, or poor, performers) with the median inter-category edit distance (across the good and poor performers; Methods, Fig. 2E). These differences were also not significant (p>0.1, one-tailed signed rank test).

**Figure 2.**
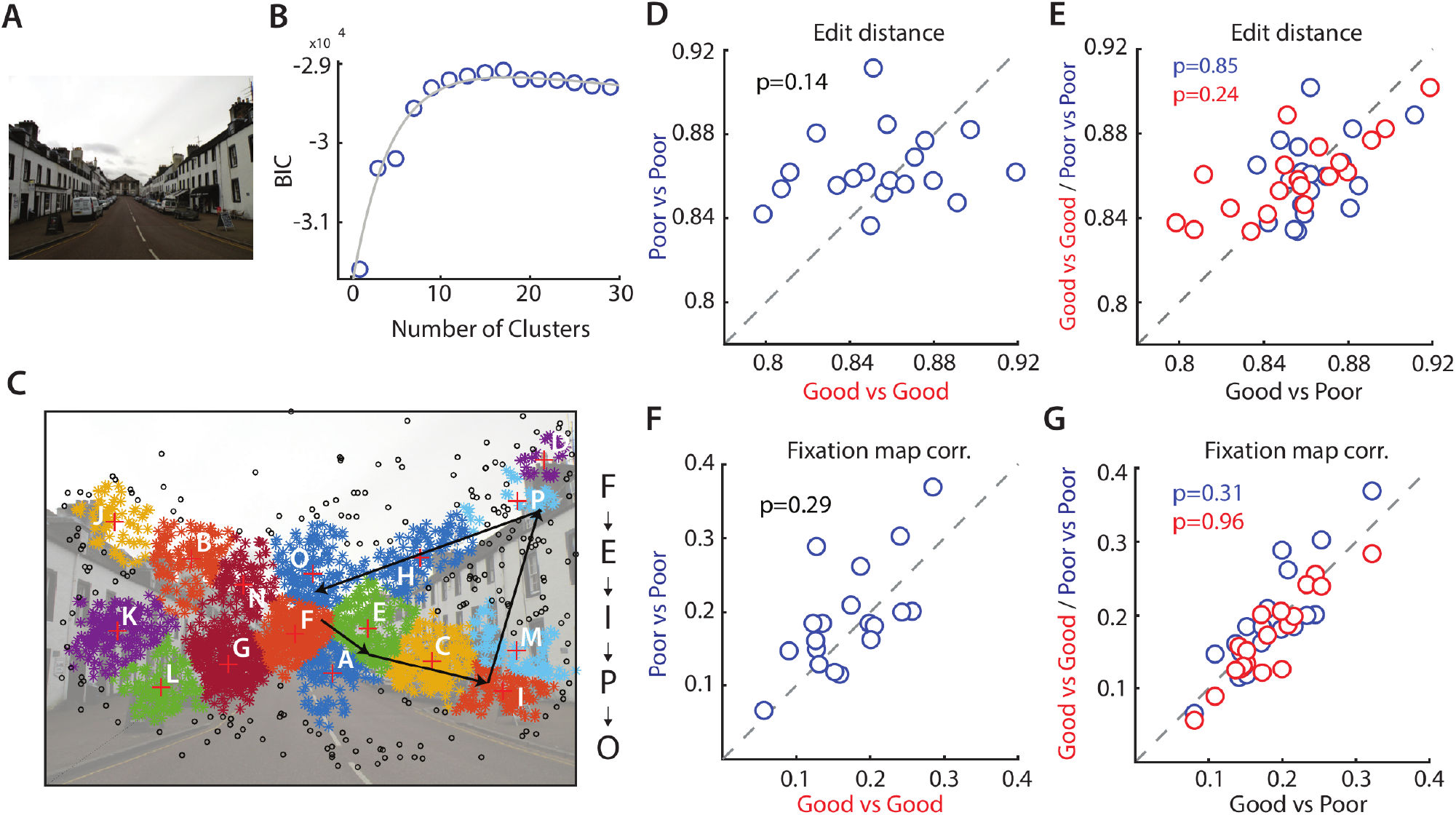
Scan paths and fixation maps do not distinguish good from poor performers. **A.** Representative image used in the change blindness experiment (Image #6, SI Fig. S1). **B.** Variation in the Bayesian Information Criterion (BIC; y-axis) with clustering fixation points into different numbers of clusters (x-axis; Methods). Circles: Data points. Gray curve: Bi-exponential fit. **C.** Clustering of the fixation points based on the peak of the fitted BIC (n=13) profile. Fixation points in different clusters are plotted in different colors. Black fixations occurred in fixation sparse regions that were not included in the clustering. Black arrows show a representative scan path – a sequence of fixation points. The character “string” representation of this scan path is denoted on the right side of the image. **D.** Distribution of edit distances among good performers (x-axis) versus edit distances among poor performers (y-axis). Each data point denotes median edit distance for each image tested (n=19). Other conventions are the same as in Figure 1E. **E.** Distribution of intra-category edit distance (y-axis), among the good or among the poor performers, versus the inter-category edit distance (x-axis), across good and poor performers. Red and blue data: intra-category edit distance for good and poor performers respectively. Each data point denotes the median for each image tested (n=19). Other conventions are the same as in panel D. **F.** Same as panel D, but comparing Pearson correlations of fixation maps among good (x-axis) and poor performers (y-axis). Other conventions are the same as in panel D. **G.** Same as in panel E, but comparing intra- versus inter-category Pearson correlations of fixation maps. Other conventions are the same as in panel E.

Second, we computed the probabilities of saccades among specific types of clusters (“domains”, Methods), which corresponded to groups of clusters ranging from most visited, on average, to least visited, for each image. We tested if the saccade probabilities among domains were different between good and poor performers. For illustration, a single saccade probability matrix, estimated by pooling scan paths across subjects is shown in Figure 3A; these correspond to inter-domain saccade probabilities for good performers (top matrix), and poor performers (bottom matrix), respectively (average across n=19 images). Visual inspection of these saccade probability matrices reveals no apparent differences. In addition, we sought to classify these saccade probability matrices, by computing them for each subject, and employed linear SVMs to classify these as belonging to good or poor performers. The SVM classification accuracy was low (~56.67%, Fig. 3B) and was not significantly different from chance (p>0.1, permutation test).

**Figure 3.**
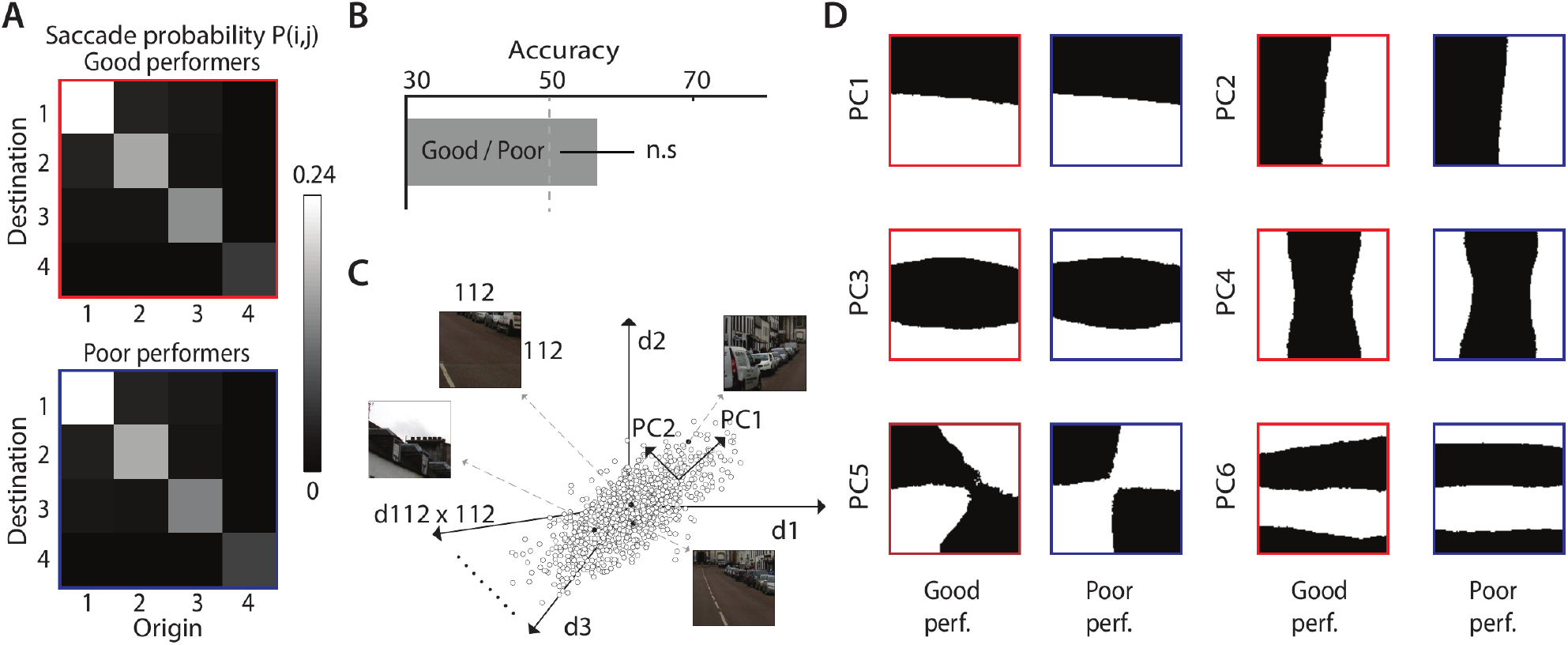
Good and poor performers exhibit similar saccade probabilities and fixated features. **A.** Average saccade probability matrices for the good performers (top; red outline) and poor performers (bottom; blue outline). These correspond to probabilities of saccading among different “domains” (1-4), each corresponding to a (non-contiguous) collection of image regions, ordered by frequency of fixations: most fixated regions (domain 1) to least fixated regions (domain 4). Cell (i, j) (row, column) of each matrix indicates the probability of saccades from domain j to domain i. **B.** Classification accuracy for classifying good versus poor performers based on the saccade probability matrix features, using a support vector machine classifier. Other conventions are as in Figure 1D. **C.** Identifying low-level fixated features across good and poor performers. 112×112 image patches were extracted, centered around each fixation, for each participant; each point in the 112×112 dimensional space represents one such image patch. Principal component analysis (PCA) was performed to identify low-level spatial features explaining maximum variance among the fixated image patches, separately for good and poor performers. **D.** Top 6 principal components, ranked by proportion of variance explained, corresponding to spatial features explaining greatest variance explained across fixations, for good performers (left panels) and poor performers (right panels). These spatial features were highly correlated across good and poor performers (median r=0.20, p<0.001, across n=150 components).

Third, we computed fixation “maps”: two-dimensional histograms of the distribution of fixations computed from the scan path for each image (Itti, Koch, and Niebur 1998). We correlated the fixation maps for each image across every pair of participants, and tested if the correlations between fixation maps were significantly different between good and poor performers (Methods). Again, we observed no significant differences between fixation map correlations of good and poor performers (Fig. 2F, p=0.29, Wilcoxon signed rank test), and no significant differences between the median intra-category fixation map correlation values and the median inter-category correlation values (Fig. 2G, p>0.1, one-tailed signed rank test).

Finally, we asked if good and poor performers differed in terms of low level spatial features in the image regions where each fixated, computed using principal components analysis (PCA, Methods).

These low-level features typically comprised horizontal or vertical edges at various spatial frequencies, and were virtually identical between good and poor performers (Fig. 3D, first six principal features for each class). We performed a correlation analysis across the top ranked 150 components that explained up to 80% of the variance in fixated image patches across both groups of subjects. Average correlations across components of identical rank were highly significant across good and poor performers (median r=0.22, p<0.001, n=150 components). We also computed these fixated features with the saliency map, computed with the frequency tuned saliency (Achanta et al. 2009), rather than with the images directly. Again, fixated features were highly correlated (median r=0.1969, p=<0.001, n=150 components, SI Fig. S3), indicating that low-level fixated features did not differ, on average, between good and poor performers.

Taken together, these analyses indicate that simple gaze metrics like fixation duration and saccade amplitudes were informative about success with change detection, whereas more complex metrics like scan paths and fixated image features failed to be as informative. In other words, our results indicate that low-level gaze metrics rather than high-level scanning strategies, determined their success with change detection.

### A model of gaze fixations for the change blindness task

Based on these empirical observations, we sought to develop a neurally-inspired model of change detection that could provide a mechanistic explanation for the trends in the human data. Briefly, our model posits that observers fixated, in turn, to different regions of the image and, at each fixation, accumulated evidence for deciding between one of two possible events: change or no change at that location. If observers detected a change, they maintained gaze fixation at the change location (per the requirements of our task). Alternatively, if no change was detected, they continued to scan the image by saccading to a new, likely, location of change.

We modeled observers’ gaze strategies using the theoretical framework of Sequential Probability Ratio Testing (SPRT), with evidence accumulation modeled as an Ornstein-Uhlenbeck process (Busemeyer and Townsend 1993). Briefly, each image was divided into a grid of non-overlapping patches, each represented by a unique neural population. Such a topographic representation of visual space occurs in various regions of the brain, including the superior colliculus, which is known to be important for controlling eye movements (Knudsen 2018) and for detecting changes (Cavanaugh and Wurtz 2004). During the fixation epochs, each neural population’s firing rate was determined by the local salience of the image patch encoded by the respective population. At each instant of time, the model gathers evidence about all regions by computing a likelihood ratio – for change versus no change – based on population firing rates and accumulates this evidence, independently for each region. If the evidence accumulated at the fixated region exceeds a threshold, a change is indicated; alternatively, the next location to saccade is determined based on the likelihood ratio landscape over the entire image. The model is discussed in detail in subsequent sections.

#### Neural representation of the image

Neurons in the brain are known to have extended visual receptive fields that can span several degrees of visual space. We, therefore, modelled neural representations by partitioning each image (*A*) and its altered version (*A’*) into a uniform grid (72×54) of equally-sized regions. While in the brain, neural receptive fields typically overlap, we did not model this overlap here, for reasons of computational efficiency (Methods). We index each region for each image as *A*_1_, *A*_2_ …, *A*_*N*_ and *A’*_1_, *A’*_2_…, *A’*_*N*_, respectively (N=3888). A saliency map was also computed for each image of each pair, which we denote as *S* and *S’*, respectively; the saliency map was computed with the frequency-tuned salient region detection method (Achanta et al. 2009); this algorithm was effective at detecting color-based changes (e.g. Image #15, SI Fig. S1). In a later section, we also employ saliency computation with a deep neural network (DeepGaze II, Kümmerer, Wallis, and Bethge 2016); the reasons for employing a deep neural network are discussed in that section.

Each region’s (*A*_*i*_) visual information was encoded by a distinct neural population. Firing activity was modeled as a Poisson process, with the average firing rate of the *i*^*th*^ population given by *λ*_*i*_. Although, conventionally, single neuron firing is modeled with a Poisson process (Rieke et al. 1997), our modeling of the neural population’s activity also as a Poisson process is plausible, under the assumption that neurons within each population fire independently of the others. In this case, the sum of independent Poisson processes is also a Poisson process. The neural firing rate for each region *λ*_*i*_ was a linear function of the average saliency of image patch falling within that region, and limited to be within a physiologically plausible range (Table 1; White et al. 2017). These were computed as: 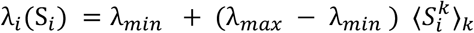, where 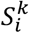 is the saliency value of the kth pixel in region *A*_*i*_, and k indexes over all the pixels in the region *S*_*i*_. In general, if the change between *A* and *A’* occurred in region *i*, the firing rates *λ*_*i*_ and *λ*’_*i*_ would differ in a manner directly related to difference in saliency across the change.

**Table 1.** Model parameters and their default values.

| Parameters | Symbol | Value | Description |
| --- | --- | --- | --- |
| Time bin | $\Delta t$ | 25 ms | Unit of time for the model |
| Image duration | $\tau$ | $10\Delta t$ | Duration for which each image or blank is shown |
| Trial duration |  | 60 s | Total duration of trial |
| Fixation interval | $q \sim N(\mu_{FD}, \sigma_{FD})$ | $\Delta t * N(15, 1)$ | Duration for maintaining fixation |
| Temperature | $T$ | 0.01 | Modulates stochasticity of next saccade |
| Decay factor | $\gamma$ | 0.004 | Decay of the evidence with time (inversely related) |
| Decay scale | $\beta$ | 4.0 grid units | Spatial range of evidence decay |
| Noise scale | $W$ | $U(-5, 5)$ | Models noise in evidence accumulation |
| Threshold | $F$ | 100 | Evidence threshold to determine change |
| Firing rate bounds | $\lambda_{min}, \lambda_{max}$ | 5, 120 spikes/bin | Minimum and maximum firing rates |
| Firing rate prior | $\mu_f$ | 3 spikes/bin | Expected difference in firing rates |

In addition, we modeled a key property of the visual system (White et al. 2017) – foveal magnification – using the cartesian variable resolution (CVR) transform (Wiebe and Basu 1997). Saliency computation was performed on the CVR-transformed image, rather than on the original image, to mimic the likely sequence of these two computations in the brain (Methods).

#### Modeling change detection

Each trial, with a total duration *T*, comprised of a large number of equal time bins of size *Δt* (Table 1). At every time bin, the number of spikes from each neural population was drawn independently of other time bins, and of other populations, from a Poisson distribution. As described above, the mean of this Poisson distribution was determined by the average saliency, within the respective region, of the CVR-transformed image. The duration of each fixation was modeled with a random variable *n*, measured in units of time-bins, such that each fixation lasted, on average *q*=*nΔt* units of time. For each fixation, *n* was drawn from a Gaussian distribution (Methods; Table 1). At the end of *q* units of time, the model either shifted gaze to a new location using a softmax strategy (described below), or confirmed having detected the change, and the simulation was terminated.

One such fixation is schematized in Figure 4A. Assume that once the fixation commences, the first image of the pair (say, *A*) persists from *t*=*1* to *t*=*m* time-bins. Following this, a blank occurs for an interval from *t*=*m*+*1* to *t*=*p* time-bins. After this, the second image of the pair (*A’*) appears for an interval from t=p+1 to t=n, until the end of fixation. We denote the number of spikes produced by neural population i at time t by 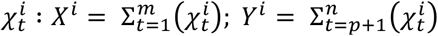; where X_i_ and Y_i_ are the total number of spikes fired by neural population i when fixating at images *A* and *A’* respectively. For convenience, we assume that no spikes occurred during the blank period 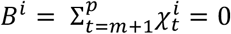, although this is not a strict requirement for the model. In general, *X*^*i*^ and *Y*^*i*^ will be statistically distinguishable if the region *A*_*i*_ of the image contains a change and vice versa.

**Figure 4.**
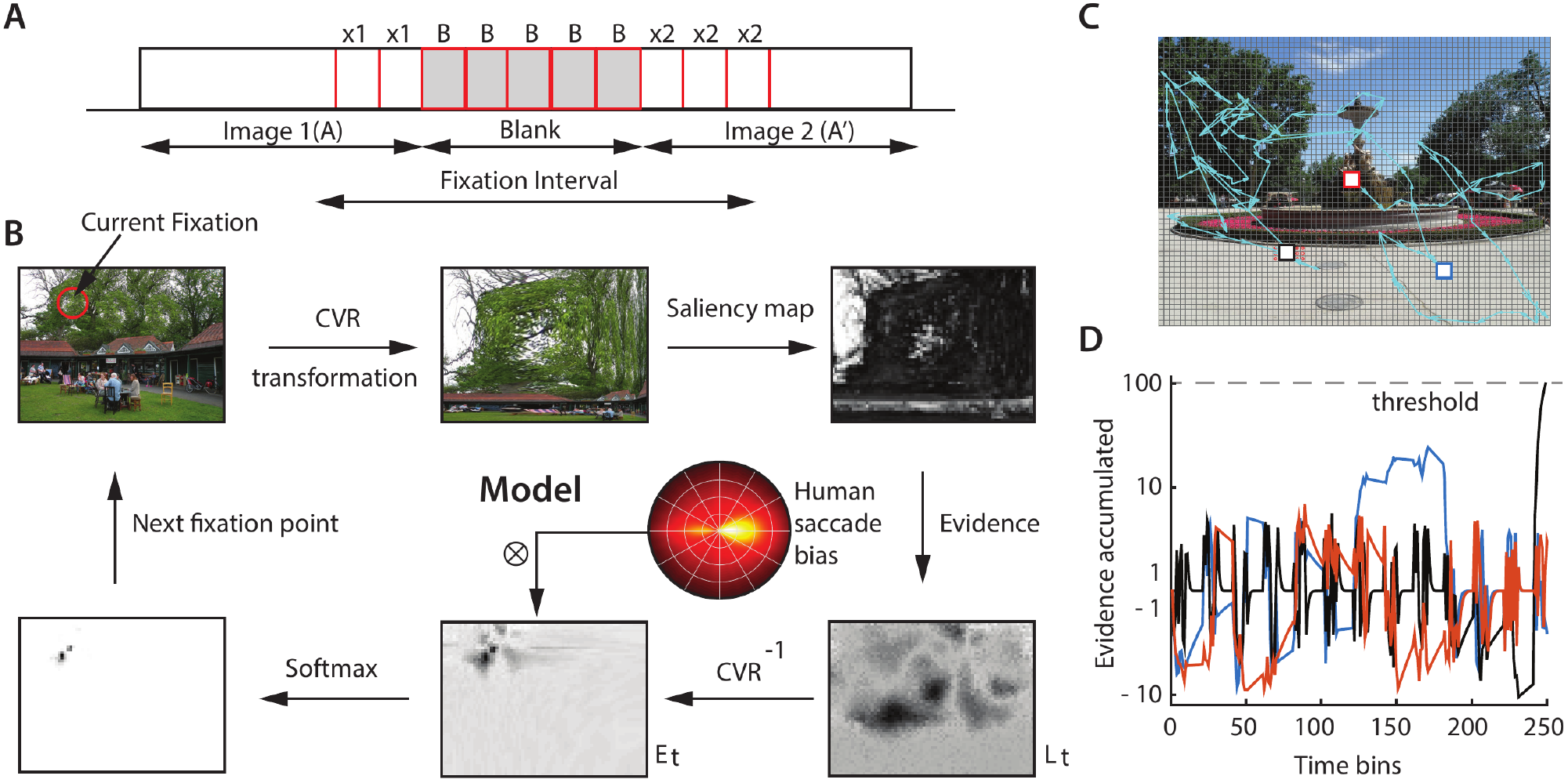
SPRT model of gaze strategies for change blindness tasks. **A.** Schematic showing a fixation across the pair of images (A, A’) and an intervening blank. **B.** Steps involved in the model’s scanning strategies (see text for details). At each point of fixation, a Cartesian variable resolution (CVR) transform is applied to mimic foveal magnification, followed by a saliency map computation to determine firing rates at each location. Instantaneous evidence for change versus no change (log-likelihood ratio, log (L(t))) is computed across all regions of the image. An inverse CVR transform is applied to project the evidence back into the original image space, where noisy evidence is accumulated, (SPRT, drift-diffusion model). The next fixation point is chosen using a softmax function applied over the accumulated evidence (E). To model human saccadic biases, a distribution of saccade amplitudes and turn angles is imposed on the evidence values prior to selecting the next fixation location. **C.** A representative gaze scan path following model simulation (blue arrows). Colored squares: specific points of fixation (see panel D). Grid: Fine divisions over which the image was sub-divided to facilitate evidence computation. Red and black squares denote first (beginning of simulation) and last (detection of change) fixation points, respectively. **D.** Evidence accumulation as a function of time for three representative regions; each color denotes evidence at the corresponding square in panel C (colored squares). When the model fixates on the red or blue squares, the accumulated evidence is indicative of no change (negative axis), or does not cross the threshold for change detection (horizontal, dashed gray line). As a result, the model continues to scan the image. When the model fixates on the black square (in the change region), the accumulated evidence crosses threshold, the change is detected, and the simulation terminates.

Nevertheless, two key challenges with change detection, both for the model and for humans, are the following: First, if the images were not interspersed by a blank, a simple differencing operation over successive time epochs (e.g. of the form *X*^*i*^ − *Y*^*i*^) may suffice to localize the change region. Yet, when images are interspersed by a blank interval, such a subtraction is not an adequate strategy: a subtraction of the form *B*^*i*^ − *X*^*i*^ or *Y*^*i*^ − *B*^*i*^ both yield large values at all locations of the image. In other words, information must be maintained over the blank interval and compared with new information following the blank, for detecting the change. Second, because the firing of Poisson neurons is stochastic, a difference in the number of spikes produced before and after the blank (*X*^*i*^ versus *Y*^*i*^) does not guarantee a change in the respective image region. Rather, such a difference may simply reflect the variance associated with the number of spikes generated by a Poisson process over these time windows. In other words, any strategy for detecting changes, must take into account the stochasticity in the neural representation of the image, both before and after the blank.

To address both of these challenges, we propose the following decision policy: the model fixates each image of the pair over an extended interval, accumulating evidence to determine whether or not a change occurred. Specifically, we model the change detection in two steps.

In the first step, the model determines, at each time-bin following the onset of the second image (*t*>*p*), whether a difference in the number of spikes (*Z*^*i*^ = *X*^*i*^ − *Y*^*i*^) between the two images at the fixated region A_i_ is attributable to Poisson variability of a single, underlying generating process (no change; N), or due to a true difference in the means of two generating processes (change; C), using a likelihood ratio rule. Assuming that the difference in the number of spikes between the two images at region *A*_*i*_ at the end of time-bin *t* is *z*, the model computes the following likelihood ratio:

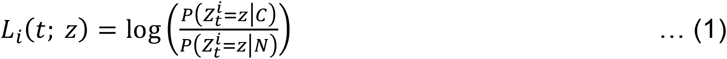

Details of computing this likelihood ratio are provided in the Methods (section on Computation of the likelihood ratio). This computation involves an infinite sum, which we calculate using Bessel functions and, in special cases, more efficiently with analytic approximations (Abramowitz and Stegun 1964). Specifically, likelihood ratio takes the form:

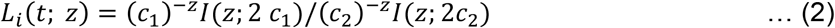

where *I*(*z*, *c*) denotes a modified Bessel function of the first kind (Abramowitz and Stegun 1964) and *c*_1_ and *c*_2_ are functions of the “expected” change in firing rates *μ*_*f*_ (prior) and the number of fixated time-bins of image and blank (*m, n, p*; Methods for details; SI Fig. S4).

In this formulation, we make the following assumptions: i) the model makes a change versus no-change decision based on the *difference* of spike counts 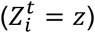, rather than the absolute spike counts produced by each image (see next); ii) the model estimates the average firing rate based on the number of spikes produced until that time-bin *λ*_*i*_ = (*X*_*i*_ + *Y*_*i*_)/(*n* − *p* + *m*); iii) the model has a discrete, single valued, *prior* on the change in firing rates *μ*_*f*_; this prior is different from the actual difference in firing rates across the two images, which is computed based on the difference in their, respective, salience values during the simulation. A later section discusses how changes in this prior affect the model’s performance (Fig. 5C).

**Figure 5.**
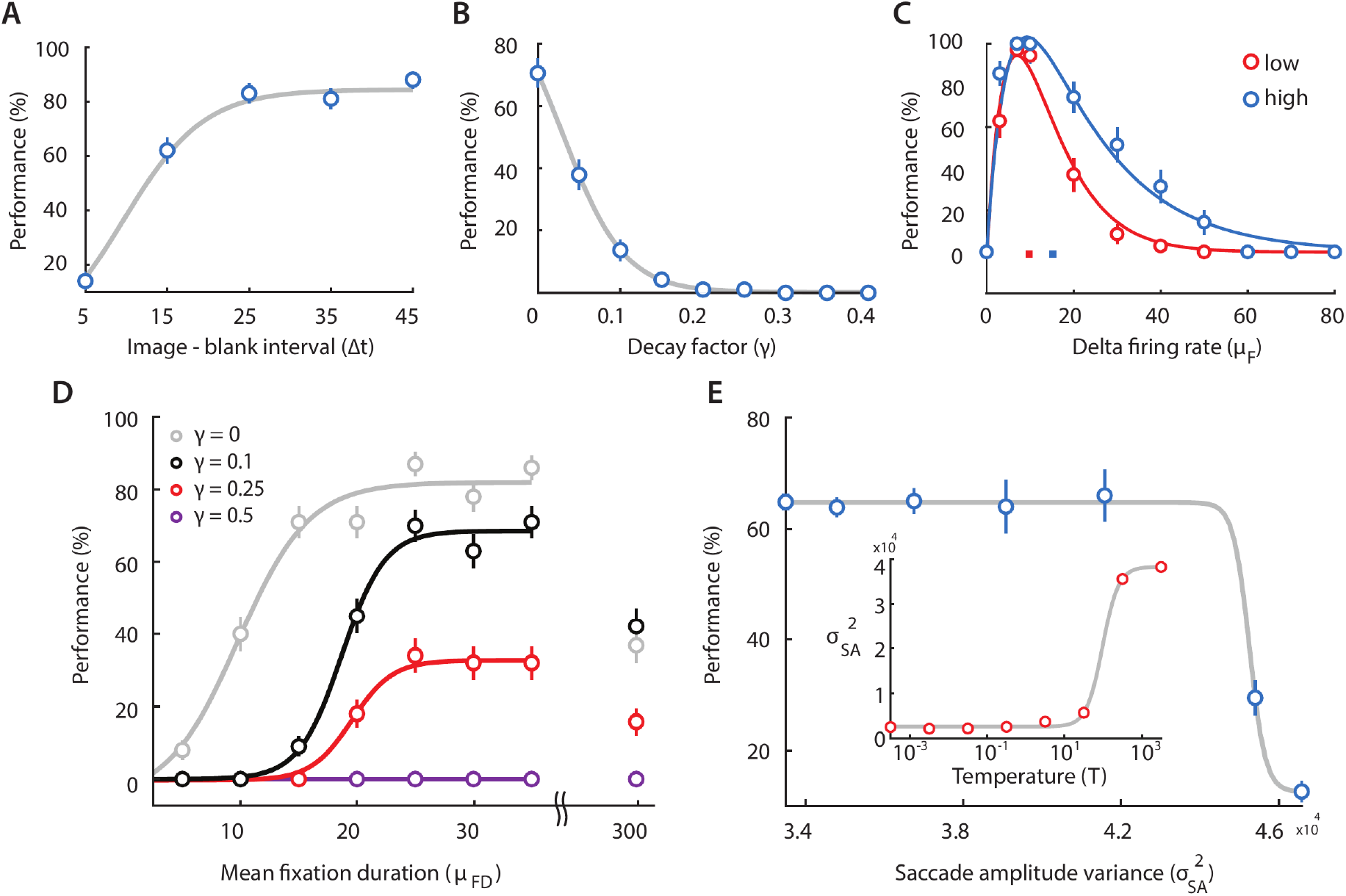
Effect of model parameters on change detection success rates. **A.** Change in model performance (success rates, % correct) with varying the relative interval of the images and blanks, measured in units of time bins (Δt=25 ms/time bin; Table 1), while keeping the total image+blank interval constant (at 50 time bins). Positive values on the x-axis denote larger image intervals, as compared to blanks, and vice versa, for negative values. Blue points: Data; gray curve: sigmoid fit. **B.** Same as in panel (A), but with varying the maximum decay factor (γ; equation 3). Curves: Sigmoid fits. **C.** Same as in panel (A) but with varying the firing rate prior (μ_f_) for image pairs with the lowest (red; bottom 33^rd^ percentile) and highest (blue; top 33^rd^ percentile) magnitudes of firing rate changes. Curves: Bi-exponential fits. **D.** Same as in panel (A), but with varying the mean fixation duration (μ_FD_; measured in time bins, Δt=25 ms/time bin). Each color denotes simulations for a different value of the decay factor (γ). Rightmost data (after axis break): Performance at very high mean fixation durations (300 time bins). Curves: Sigmoid fits. **E.** Same as in panel (A), but with varying saccade amplitude variance (σ^2^ _SA_). (Inset) Variation of σ^2^ _SA_ with the softmax function temperature parameter (T) (see text for details). Curves: Sigmoid fits. B-E. Other conventions are the same as in panel A.

In the second step, the model integrates the “evidence” for change, by accumulating *L*_*i*_*(t)*, with previously-acquired evidence until time-step t-1.

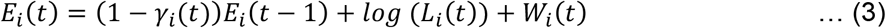

where 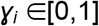 is a decay parameter for the evidence at location *A*_*i*_, which simulates “leaky” evidence accumulation (Gold and Shadlen 2007; Ratcliff et al. 2016); the larger the value of 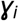, the greater the “leak” in the evidence accumulation process. The spatial distribution of the decay parameter at each region was specified based on a two-dimensional Gaussian function, with its peak at the region of fixation (Table 1); therefore, *γ*_*i*_ is a function of time and depends on the current region being fixated. In addition, at each time-bin white noise, *W*_*i*_*(t)*, sampled from a uniform distribution (interval: ±5.0, Table 1) was added to the evidence to mimic noisy evidence accumulation (Ornstein-Uhlenbeck model; Busemeyer and Townsend 1993). The integration is performed for each region in the image, independently. Both of these features – leak and noise in evidence accumulation – are routinely incorporated in models of human decision-making (Ratcliff et al. 2016), and are grounded in experimental observations in brain regions implicated in decision-making (Gold and Shadlen 2007). Also, we posit that the process of evidence accumulation occurred in the original, physical space of the image, and not in the CVR transformed space (Fig. 4B).

In summary, the model scans the image for detecting changes based on the following policy. At the beginning of each step, the model fixates at region *i* in image A for *q* time-bins (Fig. 4A). When a blank happens, and a new image appears, the model begins evaluating evidence for change. At the end of each time-bin in the second image of each pair, the model computes evidence for change (versus no change) at region A_i_ based on a likelihood ratio *L*_*i*_(*t*) (equation 1), and checks if the accumulated evidence *E*_*i*_(*t*) ≥ *F*, where *F* is the decision threshold for detecting a change (Table 1); the value of *F* controls the false-alarm (error) rate in the model. If a threshold crossing occurs, the model stops scanning the image, and region *A*_*i*_ is declared the “change region”. At the end of the fixation (*q* time-bins), if the threshold has not been crossed, the model chooses the next region *A*_*j*_ to fixate based on a soft-max probability function: 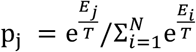, where *E*_*i*_ is the evidence value for region *i*, *N* is the number of regions in the image, and T is a temperature parameter which controls the stochasticity of the saccade (decision) policy (Methods; see also next section).

### Emergent trends in model performance resemble trends in human data

We tested the effect of key model parameters on change detection performance. These simulations provide qualitative benchmarks to assess the performance of the model against known trends in the human change blindness literature, as well as against the empirical findings in this study. These analyses were performed by using all 20 images used in the human change blindness experiment as input to the model. Default parameters were used for testing model (Table 1); detailed justification for these parameters is provided in the Methods section (Model simulation and choice of parameters). Results reported here were robust to modest variations in the values of these parameters.

Certain biases in human saccades arise from perceptual and oculomotor constraints (Roberts, Wallis, and Breakspear 2013). To account for these biases, we matched key saccade metric distributions in the human data – amplitude and turn angle of saccades – with the model (r=0.846, p<0.01); the approach for achieving this matching is described in the Methods (section on Comparison of model performance with human data). Following this, we simulated the model and measured change detection performance by varying key model parameters, including fixation duration, saccade amplitude and the like, while keeping all other parameters fixed at their default values. Results reported represent averages over several (typically 5-10) repetitions of the model over all images tested.

We illustrate the model’s policy for shifting gaze on an exemplar trial on a representative image (Fig. 4C). The model’s scan path is indicated by blue arrows showing a sequence of fixations, ultimately terminating at the change region. The figure also shows how the evidence accumulation progressed at a few, representative regions on which the model fixated (Fig. 4D, colored squares). As the model fixated on different image regions, in turn, evidence accumulated either in the positive direction, indicating potential change (Fig. 4D, e.g. blue trace, t=120-171), or in the negative direction, indicating no change (Fig. 4D, e.g. red trace, t=0-9). In each case, evidence decayed to baseline values rapidly during the blank epochs, when no new evidence was available (Fig. 4D, e.g. black trace, t=71-81). Finally, when the model fixated on the change region (black square), evidence for change continued to accumulate, until a threshold-crossing occurred. At this point, the change had been detected, and the simulation was terminated.

First, we tested the effect of varying the relative duration of the image and the blank, while keeping their overall presentation duration (image+blank) constant, on model performance. Note that no new evidence accrues during the blanks, whereas decay of accumulated evidence continues. Therefore, extending the duration of the blanks, relative to the image, should cause a substantial deterioration in the performance of the model. Indeed, this is precisely what we found (Fig. 5A): performance deteriorated (improved) systematically with decreasing (increasing) duration of the image relative to the blank. This trend mimics previously-reported patterns in human change blindness tasks, in which shortening the interval of the intervening blank improves change detection performance (Rensink, O’Regan, and Clark 1997).

Second, we tested the effect of varying the magnitude of the decay parameter (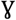, Table 1). Decreasing 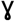 prolongs the memory for evidence relevant to change detection; 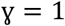 represents no memory (immediate decay; no integration) of past evidence, whereas 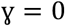 indicates reliable memory (zero decay; perfect integration) of past evidence (refer equation 3). Model success rates were near zero for 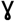 values greater than 0.4 (Fig. 5B). Following this, performance degraded systematically with increasing 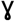. The results are consistent with neuroscience literature, which suggests that prolonging neural activity in key brain regions involved in oculomotor decisions (e.g. the superior colliculus), could facilitate performance in change blindness tasks (Cavanaugh and Wurtz 2004).

Third, we tested the effect of varying *μ*_*f*_, the prior on the magnitude of the difference between the firing rates (across the image pair) in the change region (*μ*_*Z*_, Fig. 5C). For this, we classified images into two categories – lowest and highest one-third of *μ*_*Z*_ – depending upon the range of firing rate magnitude differences for each image pair. The performance weighted average firing rate difference (*μ*_*f*_ corresponding to the center of area of the two curves) increased systematically with an increase in the prior (*μ*_*f*_). These observations suggest that the model’s ability to detect changes improves when its internal prior (expected firing rate difference) aligns with the actual firing rate difference between the image pair.

Fourth, we tested the effect of varying mean fixation duration (μ_FD_) – a key parameter identified in this study as being predictive of success with change detection. We found that increasing the mean fixation duration produced significant improvements in the performance of the model (Fig. 5D). Interestingly, after reaching a maximum, performance declined for much larger fixation duration values (Fig. 5D, values following break in x-axis). This non-monotonic trend in the model and the data can be explained as follows: Fixations which are too brief prevent accumulating sufficient evidence about the change at each region to make a reliable decision, whereas overly long fixations prevent adequate exploration of the image. There exists, therefore, an optimal fixation duration between these two extremes, governed by the magnitude of change and the time available to scan the image, which enables maximizing performance.

Finally, we tested the effect of varying the saccade amplitude variance 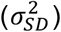 – the other key parameter we had identified as being predictive of change detection success. Because the variance of the saccade amplitude was not a parameter of the model, we varied this, indirectly, by varying the temperature (*T*) parameter in the softmax function: a higher temperature value leads to stochastic sampling uniformly from all regions of the image, whereas a low temperature value leads to more deterministic sampling, specifically from regions of maximum evidence for the change. In line with expectations, increasing *T* increased the variance of the saccade amplitudes in the model (Fig. 5E, inset). Concomitantly, with increasing saccade variance performance dropped steeply (Fig. 5E), again, recapitulating trends in the human data.

Taken together, these results show that gaze metrics, which were indicative of change detection success in humans, also systematically influenced change detection performance in the model. Specifically, the two key metrics indicative of change detection success in humans, namely, fixation duration and variance of saccade amplitude, were also predictive of change detection success in the model, suggesting a putative mechanistic link between these gaze metrics and change detection success.

### Model performance mimics human performance quantitatively

In addition to these qualitative trends, we sought to measure quantitative matches between model and human performance in this change blindness task.

First, in addition to the differences in performance across individual participants, we also observed wide variation in average detection performance, across subjects, on individual images: changes in some images were uniformly harder to detect than others (SI Fig. S2B). We tested how well correlated the model’s success rates (averaged over n=100 repetitions) were with human success rates (averaged over n=39 participants) across all images. For this analysis, we employed a saliency map generated by the DeepGaze algorithm (Kümmerer, Wallis, and Bethge 2016), rather than the frequency-tuned salient region detection algorithm (Methods). Remarkably, the model’s performance strongly correlated with human performance, across images (Fig. 6B, r=0.556, p=0.011, n=20 images). An exception was the one change image (Fig. 6B, open circle; Image #3; SI Fig. S1) that was significantly difficult to detect for humans, but not for the model (see Discussion).

**Figure 6.**
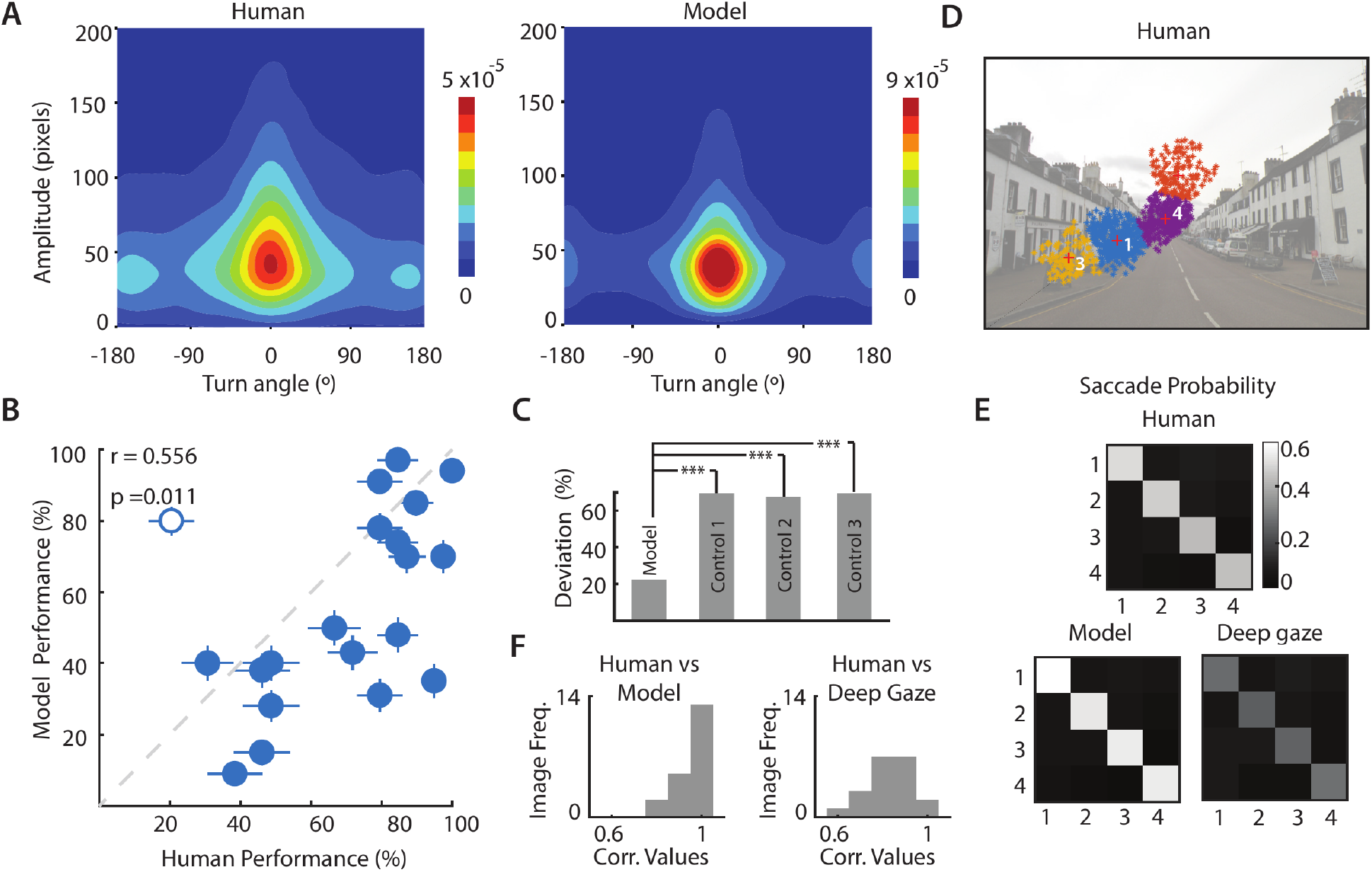
Comparison between human and model performance. **A.** (Left) Joint distribution of saccade amplitude and saccade turn angle for human participants (averaged over n=39 participants). Colorbar: Hotter colors denote higher proportions. (Right) Same as in the left panel, but for model, averaged over n=40 simulations. **B.** Correlation between change detection success rates for human participants (x-axis) and the model (y-axis). Each point denotes average success rates for each of the 20 images tested, across n=39 participants (human) or n=100 iterations (model). Open symbol: outlier image (Image #3, SI Fig. S1; see text for details). Error bars denote standard error of the mean across participants (x-axis) or simulations (y-axis). Dashed gray line: line of equality. **C.** Deviation of the model’s performance from that of humans (left bar) using the default configuration (Model), after substantially increasing the temporal decay of evidence (Control 1; γ=1), after randomly flipping the log-likelihood ratio during evidence accumulation (Control 2; ±log(L(t))) after setting the model to have a high saccade amplitude variance (Control 3; T = 10^4^). p-values denote significance levels following a paired signed rank test, across n=20 images (***p < 0.001). **D.** Top four clusters of human fixations, ranked by cumulative fixation duration, for a representative image (Image #6, SI Fig. S1). Increasing indices correspond to progressively lower cumulative fixation durations. **E.** Saccade probability matrix among the top 4 clusters with the highest cumulative fixation duration (1-4). (Top) Saccade probability matrix averaged across all images and participants. (Bottom) Saccade probability matrix for simulations of the model (left) and DeepGaze algorithm (right). **F.** Distribution, across images, of the correlations (r-values) of saccade probability matrices between human participants and the model (left) and human participants and the DeepGaze algorithm (right).

To rule out trivial reasons for these correlations, such as differences in the extent of the change region across images, we performed three different control analyses. We tested three change detection algorithms with distinct, sub-optimal search strategies: i) an algorithm that failed to accumulate evidence correctly (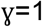; equation 3) ; ii) an algorithm in which evidence accumulation was intact, but that failed to correctly compute evidence (*log(Li(t))*; equation 1); and iii) an algorithm that accurately computed and integrated evidence but employed a random search strategy (high temperature, *T*=*10*^*4*^, in the softmax function). For each of these models, the average absolute difference in performance with the human data was significantly higher, compared with that of the original model (Fig. 6C; p<0.01, Wilcoxon signed-rank test across n=20 images).

Finally, we tested whether model gaze strategies could predict human gaze strategies beyond what was predicted by state-of-the-art fixation prediction algorithms. For this, we identified the top 4 clusters with the largest number of fixation (e.g. Fig. 6D) and computed the probability of saccades between these clusters in the human data. We compared these human saccade probability matrices with those derived from simulating the model with DeepGaze used for saliency computation (Fig. 6E, lower left).

We observed strong correlation between the human saccade probabilities and our model’s saccade probabilities, among the same fixation clusters (median r=0.94, across images; Fig. 6F, left). Next, we compared human saccade probabilities with those derived by simulating the DeepGaze algorithm *as is* without computing evidence for change (Fig. 6E, lower right; Methods). We observed considerably weaker correlations between the human data and the DeepGaze algorithm saccade probabilities (median r=0.82; Fig. 6F, right), and this correlation was significantly lower than that with the model (p<0.01 for significant difference in correlation values, signrank test).

In summary, success with change detection was robustly correlated between human participants and the model. Moreover, our model outperformed a state-of-the-art algorithm in predicting gaze shifts among the most probable locations of human gaze fixations in this change blindness task. Taken together, these findings indicate that our model provides key mechanistic insights into human gaze strategies in change blindness tasks.

## Discussion

The phenomenon of change blindness reveals a remarkable property of visual processing in the brain: Despite the apparent richness of visual perception, the brain encodes our visual world extremely sparsely. Stimuli at locations to which attention is not explicitly directed are are not effectively processed (Rensink, O’Regan, and Clark 1997). Even salient changes in the visual scene fail to be detected if these fail to capture attention. Visual attention, therefore, plays a critical role in deciding the nature and content of information that is encoded sparsely by the visual system. Gaze is typically directed to the locus of overt visual attention, and research, spanning several decades, indicates an intricate link between the neural machinery directing eye-movements and spatial attention (Gold and Shadlen 2007). In this study, we asked, therefore, if attentional strategies underlying change detection success could be understood by studying human observers’ gaze metrics and eye movement strategies in a change blindness task. Our results indicate that two low-level gaze metrics – average fixation duration and the variance of saccade amplitudes – sufficed to distinguish highly successful participants (good performers) from participants with comparatively lower success rates (poor performers). Specifically, the average fixation duration was significantly higher in good performers (Fig. 1E) and the variance of saccade amplitude was lower (Fig. 1F). These results lead to the testable hypothesis that spatial attention shifts more slowly in time, and more uniformly (or less erratically) in space, in participants who were able to detect changes effectively.

We next asked how these specific gaze metrics could explain improved success with change detection in our change blindness task. For this, we developed, *ab initio*, a neurally-constrained model based on the Bayesian framework of sequential probability ratio testing (Gold and Shadlen 2007; Ratcliff et al. 2016). The model computed evidence for change based on the likelihood ratio at each time step, and a change was detected when the evidence integrated, over time, reached a threshold (or bound). Such biologically plausible evidence accumulation (diffusion-decision) models have been widely employed in the analysis of human behavior (Ratcliff et al. 2016), and have also found correlates in neural data recorded from key brain areas, including the parietal cortex, involved in behavioral decisions (Gold and Shadlen 2007). In addition, we incorporated key neural constraints in the model’s evidence computation, including stimulus encoding with Poisson statistics over time and foveal magnification of stimulus representations in space. Despite these constraints, our model was able to faithfully reproduce key trends in the human data, in terms of variation of change detection success with fixation duration and saccade amplitudes. The results suggest a putative mechanistic link between human gaze metrics and change detection success.

Our study follows a rich literature on human gaze models, that fall, loosely, into two classes. The first class of “static” models use information in visual saliency maps (Hoffman and Subramaniam 1995; Itti, Koch, and Niebur 1998; Judd et al. 2009) to predict gaze fixations. These saliency models, however, do not capture dynamic parameters of human eye fixations, which are important for understanding strategies underlying visual exploration in search tasks, like change blindness tasks. The second class of “dynamic” models seek to predict the temporal sequence of gaze shifts. For example, Stark et al. (Hacisalihzade, Stark, and Allen 1992) proposed a model to predict saccade sequences based on finite Markov chains, whereas Wang et. al (Wang et al. 2011) proposed a model based on the principle of information maximization, whereas other models have relied on random walks and foraging techniques (Boccignone 2015; Boccignone and Ferraro 2004) and some have incorporated biological constraints also (Adeli, Vitu, and Zelinsky 2017). In a hybrid approach, Peters and Itti (Peters and Itti 2007) used an interactive visual task (playing contemporary video games) and showed that a combined model incorporating both bottom-up saliency obtained with the Itti-Koch saliency algorithm and top-down task relevance obtained using the recorded eye fixations of the human observers predicts fixations with higher accuracy than either of the approaches alone. Nevertheless, these approaches were developed for free-viewing paradigms, and comparatively few studies have focused on gaze sequence prediction during search tasks (Boccignone, 2015; Boccignone & Ferraro, 2004). To the best of our knowledge, no models have been developed for modeling gaze patterns, specifically, in change blindness tasks.

In our study, participants exhibited a surprising degree of variability with detecting changes: interindividual success rates varied across a two-fold range (Fig. 1B). These differences in performance do not necessarily reflect stable “traits” indexing participants’ inherent change detection abilities and could have been influenced by several factors, including their level of motivation and alertness during the experimental session. Nevertheless, of particular interest, is a recent study on change (Andermane et al. 2019) that evaluated test-retest reliability in change blindness tasks. This study found that observers’ change detection performance generalized over several standard tasks, and was relative stable over periods of 1-4 weeks. Moreover, using a battery of additional tasks that tested visual short term memory and attention capacities, this study identified two factors that were critical for predicting change detection success: “visual stability” – the ability to form stable and robust visual representation – and “visual ability” – indexing the ability to robustly maintain information in visual short term memory. Other studies have identified associated psychophysical factors, including attentional breadth (Pringle et al. 2001), visual memory (Angelone and Severino 2008) and familiarity with context of the visual stimulus (Werner and Thies 2000), as being predictive of change detection success.

The psychophysical factors identified by these studies have interesting parallels with key parameters in our model (Table 1). We speculate that model parameters such as the length of fixation durations (μ_FD_) and variability of saccade amplitudes (T, σ_SA_) may correspond to the “visual stability” factor. In contrast, other parameters, such as the temporal decay factor (γ) and spatial decay scale (β) may correspond to visual memory and attentional breadth variables, respectively; these latter variables may also constitute components of the “visual ability” factor identified by (Andermane et al. 2019). Finally, the firing rate prior (μ_f_) may reflect familiarity with the context of the visual stimulus. Future studies – that combine eye-tracking with behavioral assessment of these factors, among others (Smith and Milne 2009) – will be necessary to confirm these relationships.

In addition to predicting qualitative trends in the data, our model’s performance also quantitatively correlated with human performance across almost all images tested (Fig. 6B). An interesting exception to these predictions was the model performance on image #3 (SI Fig. S1A). The change in this image involved the disappearance and reappearance of a single flower in a field of flowers, which humans found quite difficult to detect, on average (20.5±6.6% success), compared to the model (80.0±4.0% success). The difference can be attributed to the perceptual phenomenon of “crowding” (Simons and Ambinder 2005; Simons and Rensink 2005), which renders it challenging for human observers to identify or recognize objects flanked by other similar or identical objects. Clearly, such detrimental effects of crowding were absent in our model. An alternative explanation for this discrepancy is that humans found the change difficult to detect due to its low “semantic” saliency. A previous report (Stirk and Underwood 2007) found that change detection success (and speed) were not significantly influenced by low-level visual saliency of stimuli. Rather, change detection success was higher when the change occurred in objects that were scene-inconsistent or semantically salient, and vice versa. Incorporating semantic saliency into the saliency map computation in our model could address this difference with the human data. Another key limitation of the model is that fixation durations were sampled from a distribution determined apriori (Table 1). Rather, future versions of the model could incorporate a likelihood ratio based rule to break fixations, for example, when the evidence against a change at that location reaches a particular threshold.

Nevertheless, the model revealed other key trends that matched human data. First, it accurately predicted human saccade data, in terms of the probability of saccades among the most likely locations of human gaze fixations. In this prediction it outperformed a model based on a state-of-the-art fixation prediction algorithm (DeepGaze; Kümmerer, Wallis, and Bethge 2016 ; Fig. 6E), indicating that the computation of the likelihood ratio for predicting the change was relevant to the model’s performance matching that of human participants. Second, the model also revealed, as an emergent feature, a common phenomenon in computational models of visual search: inhibition of return (Najemnik and Geisler 2005). This refers to the tendency of human observers to avoid visiting recently-fixated regions of visual space, and is often explicitly modeled as an additional feature in previous models (Adeli, Vitu, and Zelinsky 2017; Najemnik and Geisler 2005). Such inhibition of return is an emergent feature of our model because the accumulated evidence against a change at each location decays gradually (based on the decay factor; e.g. Fig. 4D, t=71-81, black trace), and prevents the model from fixating again at that location for several time steps.

An emerging frontier of research seeks to develop low-power autonomous agents that can be deployed independently in real-world environments. For this application, neuromorphic chips, which use spikes for visual computations, have been recently developed (Esser et al. 2016; Merolla et al. 2014). Our model is also relevant for such neuromorphic applications with spiking networks. Consider an agent surveying an environment where visual input is noisy. This noise could occur either because of poor resolution of the sensors or because of the stochastic (e.g. Poisson) spiking in neuromorphic hardware. Visual input to the agent could be interrupted (blanked) either by design or by obstacles that occur in the agent’s field of view. The agent must still detect important change events in the scene for further analysis (e.g. zooming in) or decision making (e.g. moving away). Our model provides an effective recipe for detecting changes, even with noisy inputs: the likelihood ratio (LR)-based evidence integration rule works robustly, and analytic approximations (equation 2) permit efficient LR computations. Although the computation involving Bessel functions may seem difficult to realize in neuromorphic hardware, as with neural computations in the brain, the resultant values (SI Fig. S4) are readily approximated with simple piece-wise linear functional forms; the latter are readily generated by neurons in hardware, and in the brain (Dayan and Abbott 2005; Heiberg et al. 2018; Izhikevich 2007). Implementations of the model in hardware agents may provide a test bed for investigating mechanistic nature of attentional lapses, which occur during change blindness in real-world scenarios.

## Materials and Methods

### Ethics declaration

Informed consent was obtained from all participants, and experimental protocols were approved by the Institute of Human Ethics Review board at the Indian Institute of Science (IISc), Bangalore.

### Experimental protocol

We collected data from n=44 participants (20 females; age range 18-55 yrs) with normal or corrected-to-normal vision. Of these, data from 4 participants, who were unable to complete the task due to fatigue or physical discomfort, were excluded. Data from one additional participant was irretrievably lost due to logistical errors. Thus, we analyzed data from 39 participants (18 females).

Images were displayed on a 19-inch Dell monitor at 1024×768 resolution. Subjects were seated, with their chin and forehead resting on a chin rest, with eyes positioned roughly 60 cm from the screen. Each trial began when subjects continually fixated on a central cross for 3 seconds. This was followed by presentation of the change image pair sequence for 60s: each frame (image and blank) was 250 ms in duration. The trial persisted until the subjects indicated the change by fixating at the change region for at least 3 seconds continuously (“hit”), or if the maximum trial duration (60s) elapsed and the subjects failed to detect the change (“miss”). An online algorithm tracked, in real-time, the location of the subjects’ gaze and signaled the completion of a trial based on whether they were able to fixate stably at the location of change; for each image, the areas in which a 3s stable fixation would signal a “hit” are marked in yellow in SI Figure S1. Each subject was tested on either 26 or 27 image pairs, of which 20 pairs differed in a key detail (SI Fig. 1); we call these the “change” image pairs. The remaining image pairs (7 pairs for 30 subjects and 6 pairs for 9 subjects) contained no changes (“catch” image pairs); data from these image pairs were not analyzed for this study (except for computing false alarm rates, see next). To avoid biases in performance, the ratio of “change” to “catch” trials was not indicated to subjects beforehand, but subjects were made aware of the possibility of catch trials in the experiment. We employed a custom set of images, rather than a standardized set (e.g. Rensink, O’Regan, and Clark 1997), due to the possibility that some subjects were familiar with change images used in earlier studies.

Overall, the proportion of false-alarms – proportion of fixations with durations longer than 3s in catch trials – was negligible (~0.06%, 17/32248 fixations across 264 catch trials) in this experiment. To further confirm if the subjects indeed detected the change on hit trials, a post-session interview was conducted in which each subject was presented with one of each pair of change images in sequence and asked to indicate the location of perceived change. The post-session interview indicated that about 5.7% (31/542) of hit trials were not recorded as such; in these cases, the total trial duration was 60 s indicating that even though the subject fixated on the change region, the online algorithm failed to register the trial as a hit. In addition, 2.9% (7/238) of miss trials, in which the subjects were unable to detect the location of change in the post-session interview, ended before the full trial duration (60 s) had elapsed; in these cases, we expect that subjects triggered the termination of the trial by accidentally fixating for a prolonged duration near the change. We repeated the analyses excluding these 4.8% (38/780) trials and observed results closely similar to those reported in the text. Finally, eye-tracking data from 0.64% (5/780) trials were corrupted and, therefore, excluded from all analyses.

Subjects’ gaze was tracked throughout each trial with an iViewX Hi-speed eye-tracker (SensoMotoric Instruments Inc.) with a sampling rate of 500 Hz. The eye-tracker was calibrated for each subject before the start of the experimental session. Various gaze parameters, including saccade amplitude, saccade locations, fixation locations, fixation durations, pupil size, saccade peak speed and saccade average speed, were recorded binocularly on each trial, and stored for offline analysis. Because human gaze is known to be highly coordinated across both eyes, only monocular gaze data was used for these analyses. Each session lasted for approximately 45 minutes, including time for instruction, eyetracker calibration and behavioral testing.

### SVM classification and feature selection based on gaze metrics

We asked if subjects’ gaze strategies would be predictive of their success with detecting changes. To answer this question, as a first step, we tested if we could classify good versus poor performers (Fig. 1C) based on their gaze metrics alone. As features for the classification analysis, we computed the mean and variance of the following four gaze metrics: saccade amplitude, fixation duration, saccade duration and saccade peak speed. We did not analyze two other gaze metrics acquired from the eyetracker: saccade average speed and pupil diameter for these analyses. Saccade average speed was highly correlated with saccade peak speed across fixations (r=0.93, p<0.001), and was a redundant feature. In addition, while pupil size is a useful measure of arousal (C.-A. Wang et al. 2018), it is often difficult to measure reliably, because slight, physical movements of the eye or head may cause apparent (spurious) changes in pupil size that can be confounded with real size changes. Before analysis, feature outliers were removed based on Matlab’s *boxplot* function, which considers values as outliers if they are greater than *q*_3_ + *w* × (*q*_3_ − *q*_1_) or less than *q*_1_ − *w* × (*q*_3_ − *q*_1_), where *q*_1_ and *q*_3_ are the 25th and 75th percentiles of the data, respectively, and setting *w* =1.5 provides 99.3 percentile coverage for normally distributed data.

Following outlier removal, these eight measures were employed as features in a classifier based on support vector machines (SVM) to classify good from poor performers (*fitcsvm* function in Matlab). The SVM employed a polynomial kernel, and other hyperparameters were set using hyperparameter optimization (*OptimizeHyperParameters* option in Matlab). Features from each image were included as independent data points in feature space. Classifier performance was assessed with 5-fold cross validation, and quantified with the area-under-the-curve (AUC; Bradley 1997). For these analyses, we included gaze data from all but one image (SI Fig. 1B, Image 20), in which every subject detected the change correctly. Significance levels (p-values) of classification accuracies were assessed with permutation testing by randomly shuffling the labels of good and poor performers across subjects 100 times and estimating a null distribution of classification accuracies; significance values correspond to the proportion of classification accuracies in the null distribution that were greater than the actual classification accuracy values. A similar procedure was used for SVM classification of trials into hits and misses except that, in this case, class labels were based on whether the trial was a hit or a miss, and permutation testing was performed by shuffling hit or miss labels across trials.

Next, we sought to identify gaze metrics that best distinguished good from poor performers. For this we employed three different scores that identified the relative importance of each feature (Fig. 1C):

i. *Fisher score* (Boser, Guyon, and Vapnik 1992) computes the “quality” of features based on their extent of overlap across classes. In a two-class scenario, Fisher Score for the *j*^*th*^ feature is defined as,

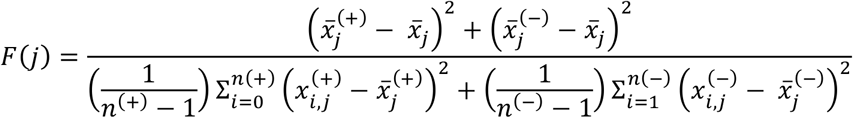

where, 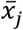 is the average value of the j^*th*^ feature. Similarly 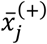 and 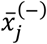 are the average of j^th^ feature for the positive and negative category respectively. Here 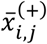 and 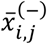 denote the j^th^ feature of i^th^ sample-index for each category, with n^(+)^ and n^(−)^ being the number of positive and negative instances respectively. A more discriminative feature has a higher Fisher score.
ii. *Sensitivity* (Nasrabadi 2007) describes the change in area-under-the-curve (AUC) with removing each feature in turn. The AUC (A) is the area under the ROC curve, plotted by varying the discrimination threshold and plotting the True Positive Rate (TPR) as a function of the False Positive Rate (FPR).

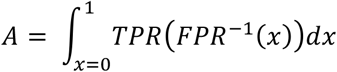

A more discriminative feature’s absence produces a higher deterioration in classification accuracy.
iii. *Information gain* (Yang 1997) is a classifier-independent measure of the change in entropy upon partitioning the data based on each feature. A more discriminative feature has a higher information gain. Given binary class labels Y for a feature X, the entropy of Y (E(Y)) is defined as,

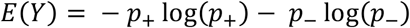

where, *p*_+_ is the fraction of positive class labels and *p*_−_ is the fraction of negative class labels.

The Information Gain given Y for a feature X is given by,

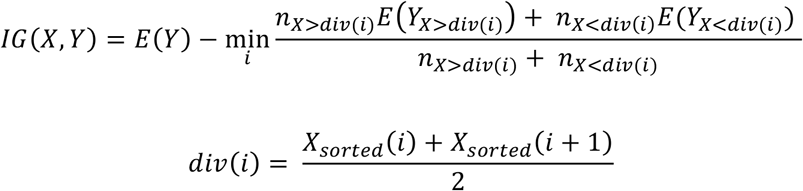

where, *n*_*X*>*div*(*i*)_ and *n*_*X*<*div*(*i*)_ is the number of entries of *X* greater than and less than *div(i)*, *Y*_*X*>*div*(*i*)_ and *Y*_*X*<*div*(*i*)_ are the entries of *Y* for which the corresponding entries of *X* are greater than and less than *div(i)* respectively and *X*_*sorted*_(*i*) indicates a feature vector with its values sorted in ascending order. A more discriminative feature has a higher Information Gain.

### Analysis of scan paths and fixated spatial features

We compared scan paths and low-level fixated (spatial) features across good and poor performers. To simplify comparing scan paths across participants, we adopted the following approach: we encoded each scan path into a finite length string. As a first step, fixation maps were generated to observe where the subjects fixated the most. Very few fixations occurred in object-sparse regions (e.g. sky), or had uniform color or texture, like the walls of a building (Fig. 2A). In contrast, many more fixations around crowded regions with more intricate details. For each image, fixation points of all subjects were clustered, and each cluster was assigned a character label. The entire scan path, comprising a sequence of fixations, was then encoded as a string of cluster labels.

Before clustering fixation points, we sought to minimize the contributions of regions with very low fixation density. To quantify this we adopted the following approach: Let *x*_*i*_ be a fixation point and let 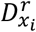 denote the average Euclidean distance of *x*_*i*_ from the set of other fixation points which are at a radius *r* from it. Let 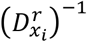 denote the inverse of 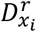. Now, we distribute all the fixation points uniformly on the image; let *U* denote this set. We find the point *y*_*i*_ in *U* that was closest (in Euclidean distance) to *x*_*i*_, and compute 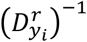, as before. Then, the fixation density at the fixation point *x*_*i*_ was defined as 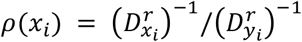. Thus, all points with density less than 1 indicate regions which were sampled with less density than that corresponding to a uniform sampling strategy. These fixation points with very low fixation density were grouped into a single cluster since these occurred in regions that were explored relatively rarely. For these analyses *r* was set to 40, although the results are robust to variations of this parameter. The remaining fixation points were clustered using k-means clustering algorithm.

The main challenges with employing the k-means algorithm were with i) deciding the number of clusters (k) and ii) deciding the initial cluster centers. For this, we employed the Bayesian Information Criterion (BIC) employed in the context of x-means clustering (Pelleg, Moore, and others 2000) to determine the optimum k. For each k ranging from 1 to 500, the BIC score was computed. Following smoothing, k corresponding to the highest BIC score was selected as the optimum cluster count. Once the number of clusters was fixed, the initial cluster centers were fixed using an iterative approach: For each iteration initial cluster centers were selected using the *k-means*++ algorithm (Arthur and Vassilvitskii 2007) and the values which gave the highest BIC score were selected as the initial cluster centers. Using the k and initial centers identified with these approaches, the fixation points were clustered for each image (Fig. 2C). Once these clusters were identified for each image, we employed two approaches for the analysis of scan-paths.

First, we computed the edit distance between scan paths (Cristino et al. 2010). For each image, the edit distance between the scan paths of each pair of subjects were calculated and normalized (divided) by the longer scan path length of the pair; this was done to normalize for differences in scan path length across subjects. A distribution of normalized edit distances was calculated among the good performers, and among the poor performers, across images. Median edit distance of each category of performers was compared against the other, with a Wilcoxon signed rank test; however, note that the lack of a significant difference would only indicate that good performers and poor performers, each, followed similarly-consistent strategies. Therefore, to test whether these strategies were indeed significantly different between good and poor performers, we compared the median edit distance among the good (or poor) performers (intra-category edit distance) with the median edit distance across good and poor performers (inter-category edit distance), for all images, with a one-tailed signed rank test.

Second, we computed the probabilities of saccading among specific types of clusters, which we call “domains”. Clusters obtained for each image were sorted in descending order of cumulative fixation duration. These were then grouped into four “domains”, based on quartiles of fixation duration, and ordered such that the first domain had the highest cumulative fixation duration (most fixated domain) and the last domain had the least cumulative fixation duration (least fixated domain). We then computed the probability of saccading from each domain to the other. We denote these saccade probabilities as: *P*(*i*_*k*_, *j*_*k+1*_), which represents the probability of saccading from domain *i* at fixation *k* to domain *j* at fixation *k+1*. We tested if the saccade probabilities among domains were different between good and poor performers by using saccade probability matrices as vectorized features in a linear SVM analysis (other details as described in section on “*SVM classification and feature selection based on gaze metrics*”).

Third, we computed the correlation between fixation distributions over images. Each image was divided into 13×18 tiles, and a two-dimensional histogram of fixations was computed for each image and participant. Binning at this resolution yielded non-empty bins for at least 15% of the bins; results reported were robust to finer spatial binning. The vectorized histogram of fixations was correlated between every pair of performers for each image, and median correlations compared across the two categories of performers, with a Wilcoxon signed rank test. As before, we also compared the median fixation correlations among the good (or poor) performers with the median fixation correlations across good and poor performers (intra- versus inter-category), for all images, with a one-tailed signed rank test.

Finally, we sought to test whether good and poor performers fixated on distinct sets of low-level spatial features in the images. For this, we identified spatial features that explained the greatest amount of variance in fixated image patches across good and poor performers. Specifically, image patches of size 112×112 pixels around each fixation point, corresponding to approximately 4° of visual angle were extracted from each image for each participant and converted to grayscale values using the *rgb2gray* function in Matlab, which converts RGB images to grayscale by eliminating the hue and saturation information while retaining the luminance. Two sets of fixated image patches was constructed separately for the good and poor performers. Each of these image patch sets was then subjected to Principal Component Analysis (PCA), using the *pca* function in Matlab, to identify low-level features in the image which occurred at the most common points of fixation across each group of subjects (Fig. 3D). We, next, sorted the PCA feature maps based on the proportion of explained variance, and correlated each pair of sorted maps across good and poor performers; in the main text, we report average correlation values across the top 150 principal component maps. We did not attempt an SVM classification analysis based on PCA features, because of the high dimensionality of the extracted PC maps (~10^4^), and the low number of data points in our experiment (~800). We also performed the same analysis after transforming each image into a grayscale saliency map using the frequency tuned salient region detection algorithm (Achanta et al. 2009). The same analyses were repeated for spatial features extracted from good and poor performers’ fixated image patches.

### Computation of the likelihood ratio (Li(t; z))

We provide here a detailed derivation of equations 1 and 2 in the main paper, involving computation of the likelihood ratio ***L***_***i***_(***t*; *z***) for change versus no change at each region *A*_*i*_. At each fixation, the model is faced with evaluating evidence for two hypotheses: change (*C*) versus no change (*N*). Note that the true difference between the firing rates of the generating processes at the change region is not known to the model, *apriori*; this corresponds to the fact that, in our experiment, the observer cannot know the precise magnitude or nature of the change occurring in each change image pair, *apriori*. We posit that the model expects to observe a firing rate difference of ±***μ***_***f***_ between the means of the two Poisson processes associated with the change region; this represents the *apriori* expectation of the magnitude of change for human observers. Here, we model this prior as a singleton value, although it is relatively straightforward to extend the model to incorporate priors drawn from a specified density function (e.g. Gaussian).

Let *X*^*i*^ and *Y*^*i*^ denote the number of spikes observed in the m and n−p time-bins that the model fixates on the two images (*A* or *A’*), respectively (Fig. 4A). Let λ_i_ denote the mean firing rate observed during this fixation, up until the current time bin; for this derivation, we posit that λ_*i*_ is measured in units of spikes per time bin; measuring λ_*i*_ in units of spikes per second simply requires multiplication by a scalar factor (Table 1), which does not impact the following derivation. The model estimates the mean firing rate over the fixation interval as *λ*_*i*_ = (*X*^*i*^ + *Y*^*i*^)/(*m* + *n* − *p*). Note that this estimate of the mean firing rate is updated during each fixation across time bins.

For hypothesis *C* to be true, *X*^*i*^ would be a sample from a Poisson Process with mean, *Γ*_1_ = *m*(*λ*_*i*_ + *μ*_*f*_) or *Γ*_1_ = *m*(*λ*_*i*_ − *μ*_*f*_) and *Y*^*i*^ would be a sample from a Poisson Process with mean, *Γ*_2_ = (*n* − *p*)(*λ*_*i*_ − *μ*_*f*_) or *Γ*_2_ = (*n* − *p*)(*λ*_*i*_ + *μ*_*f*_), respectively. Similarly, for hypothesis *N* to be true, *X*^*i*^ would be a sample from a Poisson Process with mean, *Γ*_1_ = *m*(*λ*_*i*_) and *Y*^*i*^ would be a sample from an identical Poisson Process with mean, *Γ*_2_ = (*n* − *p*)(*λ*_*i*_). For detecting changes, we assume that the model computes only the difference in the number of spikes, *Z*^*i*^ = *Y*^*i*^ − *X*^*i*^, between the two images, rather than keeping track of the precise number of spikes generated by each image. The observed difference *Z*^*i*^ could, therefore, be positive or negative.

For hypothesis C (occurrence of change), the likelihood of observing a specific value of the difference in the number of spikes across the two images *Z*^*i*^ = *z* is given as:

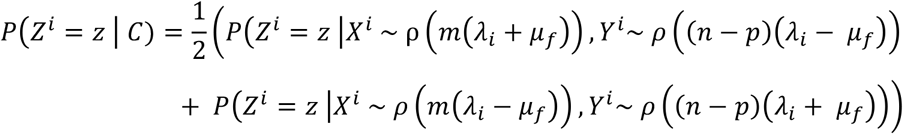

where ρ() denotes the Poisson distribution. Here, we have assumed that the prior probabilities of encountering image *A* or *A’* when the fixation lands in a given region are equal (the 1/2 factor). Similarly, for hypothesis *N* (no change), the likelihood of observing a specific difference in the number of spikes, z, is given as:

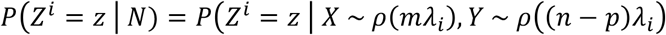

The likelihood ratio of hypotheses, change versus no change, is computed as:

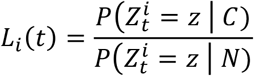

We next expand these expressions with the analytical form of the Poisson distribution, 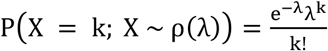, and marginalize over all values of *X*^*i*^ = *x* and *Y*^*i*^ = *x* + *z*. These calculations involve computing an infinite sum which can be efficiently solved using Bessel functions. Specifically, the infinite sum in our calculation can be computed using the identity: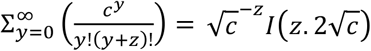 where *I* is a modified Bessel function of the first kind.

With some algebra, we can show that:

a. when 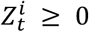:

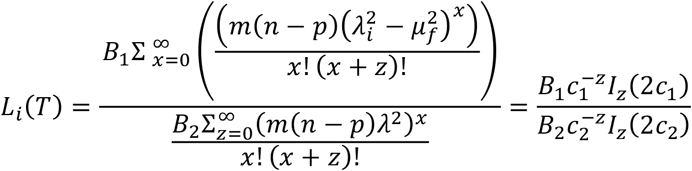

where 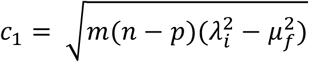 and 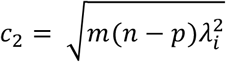, 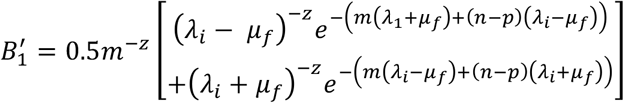 and 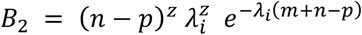.
b. when 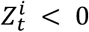:

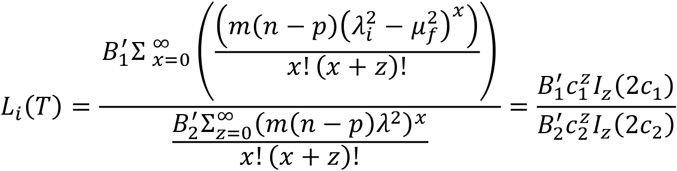

where 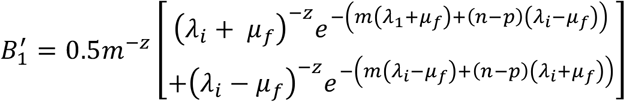,

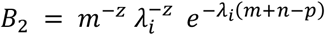

and *c*_1_ and *c*_2_ are the same as before.

We computed the value of the Bessel function using the Matlab function *BesselI*. When values of x and z were large or disproportionate, Matlab’s floating point arithmetic could not compute these expressions correctly; in this case, we employed variable precision arithmetic (*vpa* in Matlab). In addition, d for extreme values of *x* and *z* we adopted the following approximations:

a. For sufficiently large values of 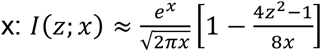
b. For sufficiently large values of 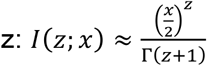

SI Figures S4A-B illustrates the variation of the logarithm of the likelihood ratio with the difference in the numbers of spikes (*z*), for different values of *λ*_*i*_ and *μ*_*f*_, respectively. Despite the detailed analysis presented in these derivations, the functional form of the log-likelihood ratio is closely approximated by a piecewise linear function which can be readily achieved by simple neural circuits (Dayan and Abbott 2005; Heiberg et al. 2018; M. Izhikevich 2007). Note that, in these calculations, we have not taken into account the prior probability of change versus no change (*P*(*C*)/*P*(*N*)). Incorporating this prior implies scaling the likelihood ratio when calculating the posterior, which produced a corresponding change in the rate of evidence accumulation (equation 3), but did not affect any of the qualitative trends reported in the results.

### Cartesian Variable Resolution (CVR) transform

We modeled a key biological feature of visual representations of images, in terms of differences between foveal and parafoveal representations. When a particular region is fixated, the representation of the fixated region, which is mapped onto the fovea, is magnified whereas the representation of the peripheral regions are correspondingly attenuated. We modeled this using the Cartesian Variable Resolution (CVR) transform, which mimics known properties of visual magnification in humans (Wiebe and Basu 1997).

The enhanced sensory representation of the foveated (fixated) region was modeled according to the following mathematical transformation of the image. We considered the foveated pixel to be the origin, denoted by (*x*_0_’ *y*_0_) in the original image. An arbitrary point in the image, denoted by (*x*, *y*) is at a distance from the origin given by, *dx* = *x* − *x*_0_ and *dy* = *y* − *y*_0_. The following logarithmic transformation was then performed:

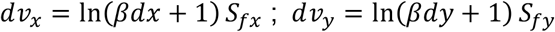

where, *β* is a constant (=0.05) that determines central magnification, and *S*_*fx*_ and *S*_*fy*_ (=200) are scaling factors along *x* and *y* directions, respectively; results reported were robust to modest variations of these parameter values. The final coordinates of the CVR transformed image are given by: *x*_1_ = *x*_0_ + *dv*_*x*_ and *y*_1_ = *y*_0_ + *dv*_*y*_.

### Model simulations and choice of parameters

The model was simulated with a sequence of operations, as shown in Figure 4B. In these simulations, the CVR transformation that mimics foveal magnification was performed before the saliency map was computed. This sequence mimics the order of operations observed in the brain: foveal magnification occurs at the level of the retina, whereas saliency computation occurs at the level of higher brain structures like the superior colliculus (Knudsen 2018) or the parietal cortex (Bisley and Goldberg 2010). Saliency maps were computed using the frequency tuned salient region detection algorithm (Achanta et al. 2009). Because of this sequence of operations, we needed to re-compute the saliency map for each image for every possible location of fixation (at the pixel level). However, this was infeasible in terms of computational time; on a standard desktop machine (Intel i7 CPU, 16 GB RAM), this computation would take an unreasonably long time (~*O(10^3^)* months). To expedite the computation, we represented each image in a reduced 864×648 pixel space and divided each image into a grid of non-overlapping patches or regions (72×54; Fig. 4B), such that each patch covered 12×12 pixels. For two images of portrait orientation (Images #10 and #19; SI Fig. S1), the same operations were done except that x- and y- grid resolutions were interchanged. We then pre-computed CVR transforms and saliency maps for each of these pre-computed grid centers and performed simulations based on these region-based representations of the images. Parcellating the image based on a finer grid, or an overlapping grid, would have improved the resolution of the saccades, but at a substantial computational cost.

Model parameters used for the simulations are specified in Table 1. The time bin (Δ*t*) was specified as 25 ms; larger and smaller values resulted in less or more frequent evaluations of the evidence (equation 1) producing correspondingly faster or slower accumulation of the evidence. The image and blank durations (τ) were fixed at 10 time bins (250 ms), matching their durations in the actual experiment. The trial duration was fixed to 2400 bins (60 s), again matching the actual experiment. The fixation interval (*q*), for each fixation, was set to be drawn from a normal distribution with mean of 15 time bins (μ_FD_=375 ms), which closely matched the median fixation duration (~300 ms) in the human data across all performers. The standard deviation of the normal distribution was set to be 1 time bin (25 ms), considerably lower than that in the human data (232 ms). A narrow distribution (with very low variance) for the fixation duration enabled robustly measuring, with only a few (5-10) simulated trials, the effect of varying the mean fixation duration on performance. The temperature (*T*) parameter was set to ensure a similar range of saccade amplitude variance in the model, as in the human data. The decay factor (γ), which determines how quickly accumulated evidence “decays” over time, and decay scale (β), which governs the spatial extent of evidence accumulation, were set to default values that enabled the model to match average human performance across all images. Then, their values were varied over a wide range to test the effect of these parameters on model success with change detection. Noise scale (*W*), which controls the noise added during the evidence accumulation process, and threshold (*F*), which controls the threshold value of evidence needed for reporting a change (Fig. 4D), were set so that their mutual values ensured negligibly low false-positive rates (< 2%), overall. Firing rate bounds (λ_min_, λ_max_) for encoding saliency were decided from biologically-plausible values for SC neurons, between 5 and 120 spikes per time bin. This corresponds to an overall population firing rate range of 0.2-4.8 kHz, which, assuming around 50 units in the neural population encoding each region, works out to a firing rate of 5-100 Hz per neuron, mimicking the biologically-observed range of firing rates for SC neurons (White et al. 2017; their Fig. 3). The firing rate prior (μ_f_) was set to 3 spikes per bin, and the effect of varying this parameter on performance was also tested (Fig. 5C). Finally, we used a third-order Taylor series approximation to the softmax function to achieve a softer saturation of this function.

Human saccade sequences tend to be biased in terms of the amplitude of individual saccades, and the angles between successive saccades (SI Fig. S5A); these biases likely reflect properties of the oculomotor system that generates these saccades (Roberts, Wallis, and Breakspear 2013). Because these saccade properties are not emergent features of our model, we matched the human saccade turn angle and amplitude distributions in the model. This was done by multiplying the map of evidence accumulated with the human saccade amplitude and turn angle distribution, before imposing the softmax function for computing saccade probabilities (Fig. 4B).

### Comparison of model performance with human data

Using the saccade generation model shown in Figure 4B, we simulated the model 100 times using the same images as employed in the human change blindness experiment (Fig. 1A). All stochastic parameters (evidence noise, fixation durations) were resampled with fresh random ‘seeds’ for each iteration of the model. We then computed the accuracy of the model as the proportion of times the model detected the change -- fixation on change region until threshold crossing (Fig. 4D) -- versus the proportion of times the model failed to detect the change region. These proportions of correct detections were then compared for human performance (average across n=39 participants) versus model performance (n=100 iterations), across images, using robust correlations (Pernet, Wilcox, and Rousselet 2013). For these analyses, we employed the state-of-the-art DeepGaze algorithm (Kümmerer, Wallis, and Bethge 2016) for generating the saliency map, rather than conventional Itti-Koch or frequency tuned salient region detection methods. DeepGaze is among the top-ranked algorithms for human gaze prediction, and is based on a deep learning model for fixation prediction which employs features extracted from the VGG-19 network, another deep learning neural network trained to identify objects in an image. In our case also, the DeepGaze algorithm, in conjunction with the model’s evidence accumulation rule, produced more robust predictions of human performance across images (Fig. 6B), rather than the frequency-tuned saliency algorithm (SI Fig. S6B). Model performance in Figure 6B (main text) was computed with μ_FD_=30, and in SI Fig. S6A with μ_FD_=15; the latter is the default value used in our simulations. Although model and human performance were significantly correlated at μ_FD_=30 (p<0.05, robust correlations) and showed a trend toward correlation at μ_FD_=15 (p=0.052, SI Fig. S2), the model performed better overall, with far fewer misses, at higher fixation durations.

Next, we performed three control analyses to ascertain that these correlations were not driven by trivial differences – for example, in the size of the change region – across images. For this we tested three different change detection models, each with a specific pattern of sub-optimality. First, we tested a model that failed to integrate evidence effectively by setting γ=1 in the evidence integration step (equation 3). Such a model completely ignores past evidence and makes decisions based solely on instantaneous evidence (*Li*(*t*)). Second, we tested a model that integrated evidence effectively, but ignored the sign (but not the magnitude) of evidence for change versus no-change at each time step. For this, we randomly multiplied *log*(*Li*(*t*)) with −1 (with 50% probability) at each time step before the likelihood ratio was incorporated into equation 3. Such a model fails to correctly take into account the evidence for change versus no change, from the likelihood ratio computed in equation 1. Finally, we tested a model in which evidence computation and accumulation were intact, but the model selected the next location of saccade with a highly random strategy. This was achieved by setting a high value of the temperature parameter (T) in the final softmax function (Fig. 4B), which resulted in a nearly uniform probability, across the image, of selecting the next fixation (“random searcher”; Najemnik and Geisler 2005). The absolute differences in performance between the human data and our model, and these control models were compared with signrank tests (Fig. 6C).

Finally, we tested model’s ability to predict human gaze shift strategies. For this we employed the following approach. First, we identified the top 4 fixated clusters in each image. Next, we constructed a saccade probability matrix between every pair of clusters among these four clusters in the human data, by pooling fixation data across all n=39 participants. The model was then simulated 100 times, and the average probability of saccades between the same clusters for each image was computed for the model. These 4×4 saccade probability matrices were then linearized and compared between the model and human data using Pearson’s correlations (Fig. 6F, left). We also tested whether this predictive power was indeed a property of the model, or was inherited from DeepGaze’s powerful saliency prediction algorithm. For this, the model was simulated with all of the same steps as in Figure 4B, except that no likelihood ratio was computed, and no evidence accumulated. In other words, saccades occurred based on DeepGaze saliency predictions alone. Because in this case the model was not accumulating evidence for change, there was no clear termination criterion, and we simulated the model for the same median number of saccades for each image, as in the original simulation that did accumulate evidence for change (Fig. 6F, right). Again, 4×4 saccade probability matrices were compared between the DeepGaze prediction and human data using Pearson’s correlations.

## Acknowledgments

We would like to thank Jogendra Nath Kundu and Neethu PM for assistance with the analyses, Ranit Sengupta for help with writing the manuscript and Guruprasath Gurusamy for help with preparing figures. This research was funded by a Wellcome Trust-Department of Biotechnology India Alliance Intermediate fellowship, a Science and Engineering Research Board Early Career award, a Pratiksha Trust Young Investigator award, a Department of Biotechnology-Indian Institute of Science Partnership Program grant, a Sonata Software foundation grant and a Tata Trusts grant (to DS).

