## Supplemental Information for "Neurally-constrained modeling of human gaze strategies in a change blindness task"

### Supplementary Information

#### Supplementary Table

##### SI Table S1. List of images employed in the change blindness task

| ID | Image | Type of Change |
|---|---|---|
| 01 | Engine Room | Appearance/ Disappearance |
| 02 | Fruitseller | Change in Size |
| 03 | Garden Ed | Appearance/ Disappearance |
| 04 | Holyrood | Appearance/ Disappearance |
| 05 | Illiterati | Appearance/ Disappearance |
| 06 | Inveraray | Change in Shape |
| 07 | Madrid | Appearance/ Disappearance |
| 08 | Market Venice | Appearance/ Disappearance |
| 09 | Meadows | Appearance/ Disappearance |
| 10 | Park Sale | Appearance/ Disappearance |
| 11 | Outdoor Party | Appearance/ Disappearance |
| 12 | Pavilion Café | Appearance/ Disappearance |
| 13 | Plaza Mayor | Change in Shape |
| 14 | Retiro Park | Appearance/ Disappearance |
| 15 | Ride Zone | Change in Color |
| 16 | Souvenir Shop | Change in Color |
| 17 | Jewellery | Change in Color |
| 18 | Visitors | Change in Color |
| 19 | Vintage Sale | Appearance/ Disappearance |
| 20 | Vatican City | Appearance/ Disappearance |

#### Supplementary Figures

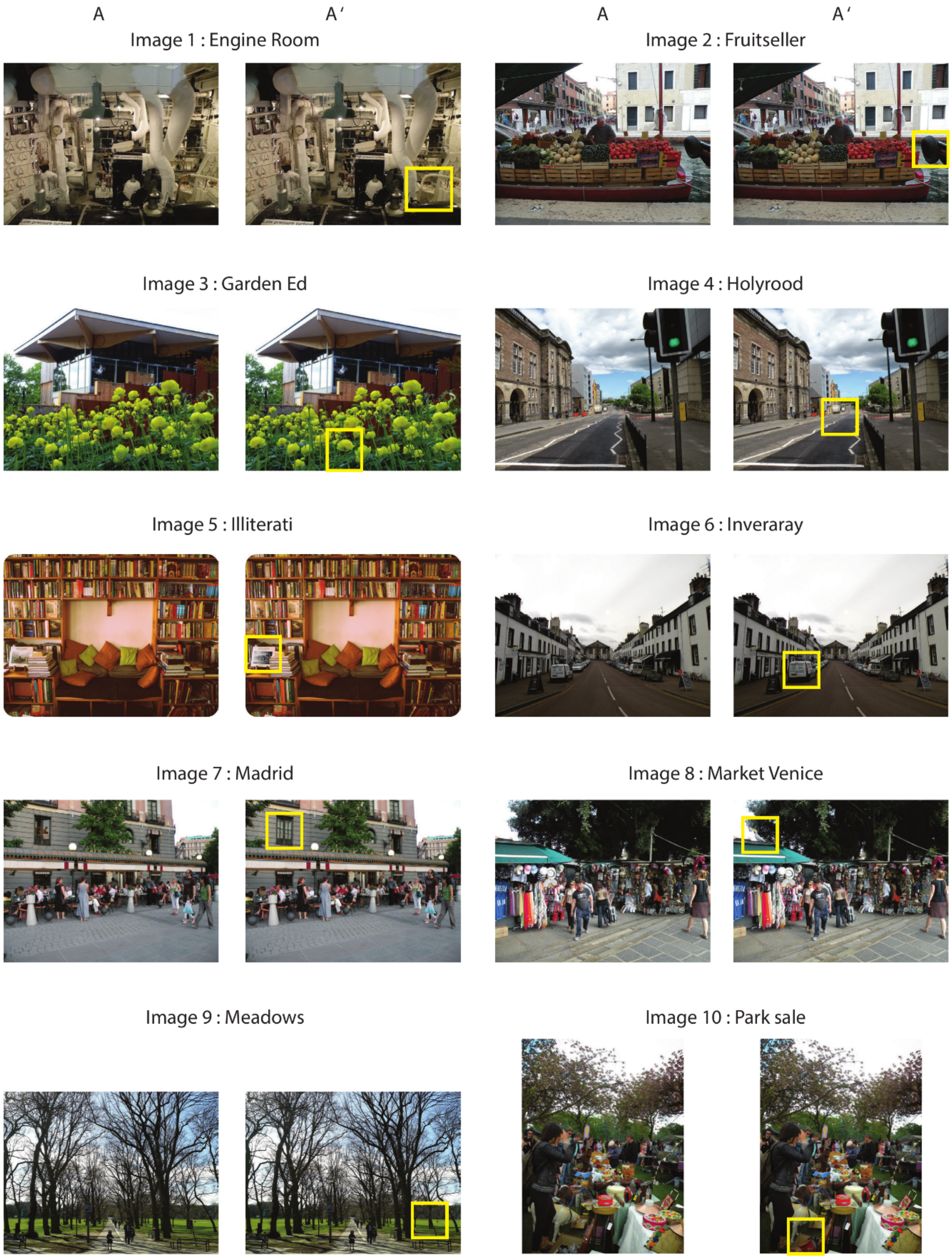


***Figure S1. List of figures used in the change blindness experiment.*** *Each sub-panel depicts the original image A (left) and its changed version A'(right). The highlighted circle indicates the location of change. This list excludes the seven catch trial images (with no change) that were included in experiments with human subjects.*

*See also SI Table S1.*

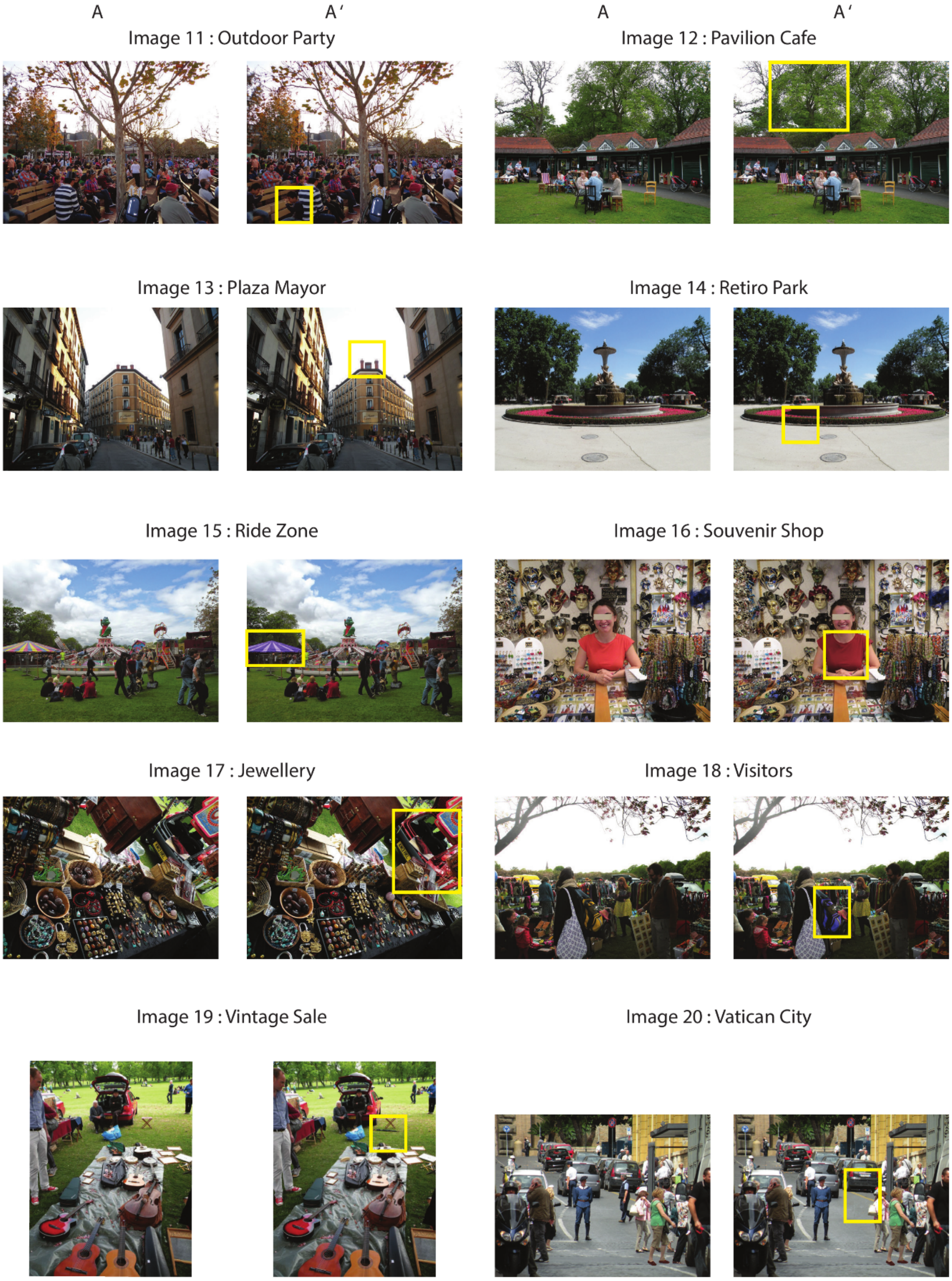


***Figure S1. List of figures used in the change blindness experiment. (cont.)***

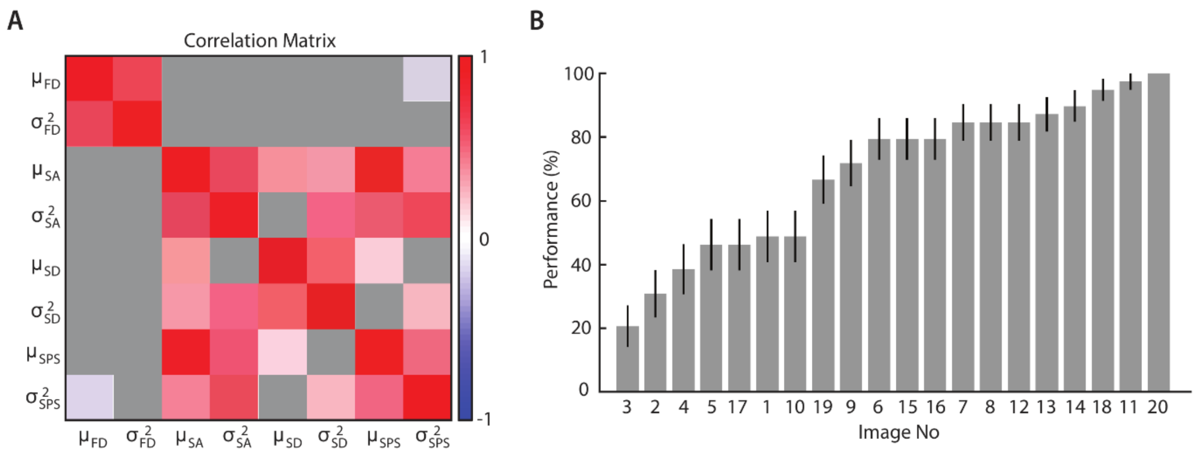


***Figure S2. Variance in success rates across images and gaze metrics predictive of success. A**. Pair-wise correlations among the eight gaze metrics used as features in classification analysis of good versus poor performers (Fig. 1C, main text). Gray squares: non-significant correlations. Colored square: significant correlations at p<0.01 with Bonferroni correction for multiple comparisons. Abbreviations are as in Fig. 1C (main text). **B**. Success rates of human observers on the change blindness trial images (n=20), sorted by the proportion of hits. Error bars denote standard error of the mean performance across participants.*

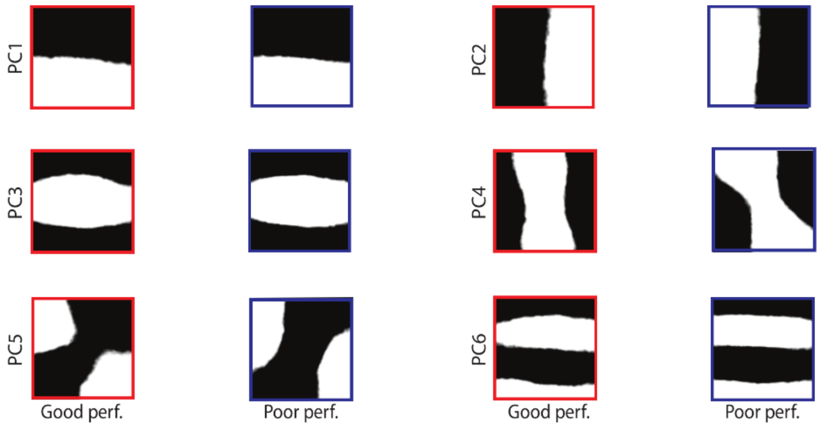


***Figure S3. Fixated features for good and poor performers identified with saliency maps.** Same as in Figure 3D (main text) except that fixated features were identified following PCA on 112x112 patches extracted from a saliency map, rather than the grayscale image. The saliency map was generated with the frequency tuned saliency algorithm (Achanta et al. 2009). Other conventions are the same as in Fig. 3D main text.*

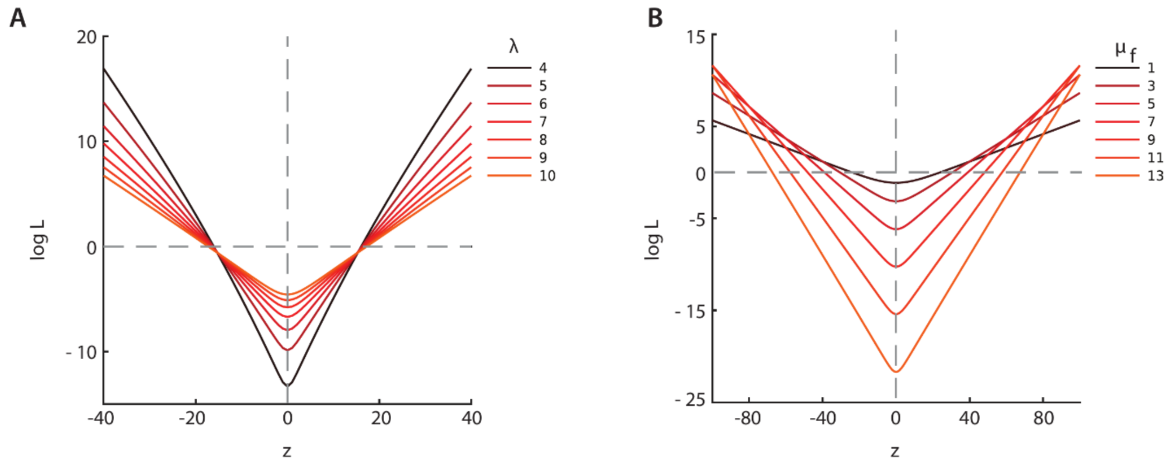


***Figure S4. Dependence of the likelihood ratio (L(t; z)) on mean firing rate and firing rate prior. A.*** *Likelihood ratio (L(t; z)) as a function of spike count difference between the first and second image (z, equation 2) for different values of the mean firing rate, λ = 4 … 10 spikes/bin. The number of time bins for which the first and second images were fixated (m and n−p, respectively) have each been fixed to 5 bins, and the firing rate difference prior,* $\mu_f$ *fixed at 3 spikes/bin. Curves of progressively lighter shades: increasing values of the mean firing rate.* ***B.*** *Same as in A, but for different values of the firing rate difference prior,* $\mu_f$ *= 1, 3, 5 … 13 spikes/bin and mean firing rate λ fixed at 40 spikes/bin. Curves of progressively lighter shades: increasing values of* $\mu_f$*.*

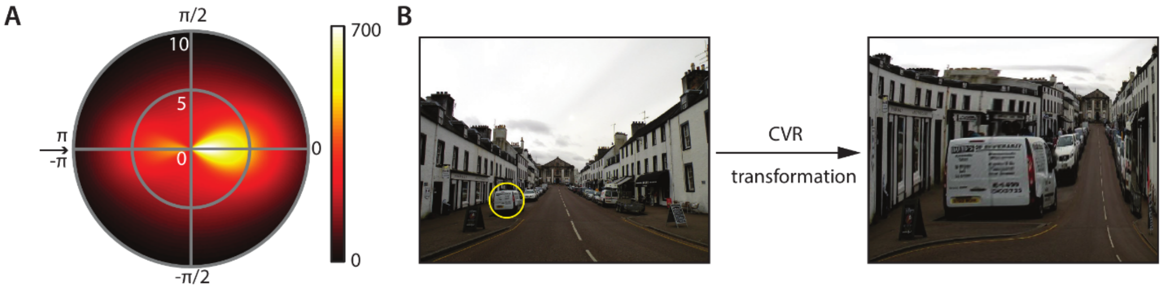


***Figure S5 Mimicking biological constraints in the model. A****. Polar heat map indicating the distribution of human saccade amplitudes and turn angles. The arrow indicates the location of the last saccade. The histogram was computed using data from all (n=39) participants and all (n=20) images. The bias against right angled turns is apparent. The distribution was smoothed both along the radial and angular directions, for display purposes only.* ***B****. Illustration of foveal magnification with the Cartesian Variable Resolution (CVR) transform for a hypothetical fixation (highlighted by the circle) on one of the images used in the change blindness task (Image #6, SI Fig. S1).*

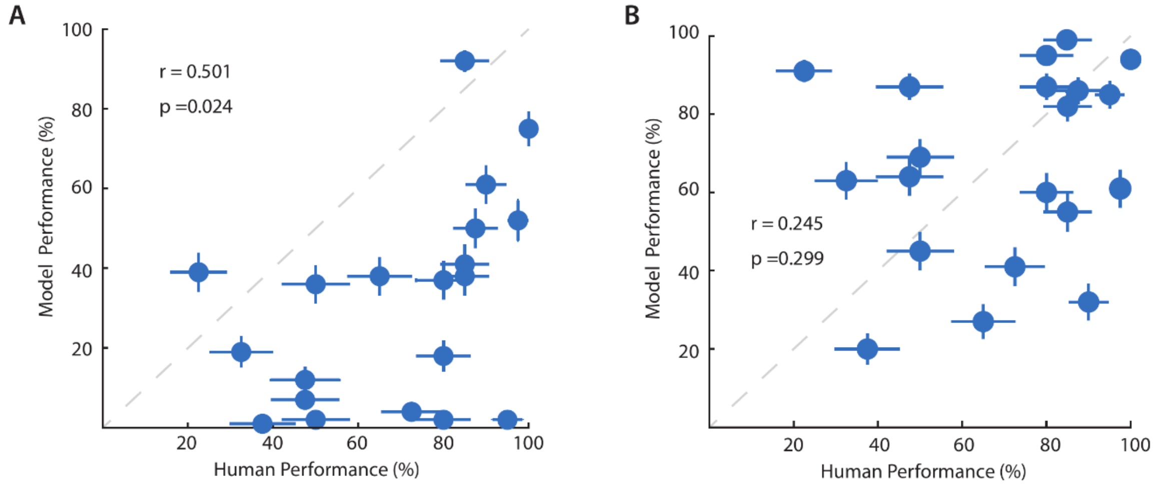


***Figure S6. Correlation between model and human performance. A.*** *Same as in Figure 6B (main text) except that the mean fixation duration was set to 15 time-bins. Other conventions are the same as in Fig. 6B (main text).* ***B.*** *Same as in panel A except that the frequency-tuned saliency algorithm was used for computing the saliency map. Other conventions are the same as in Fig. 6B (main text).*

#### Supporting Videos

*Two change blindness movies based on images 04 and 05 are provided as demos (SI Video 1 Holyrood.mp4 and SI Video 2 Illiterati.mp4). The movies show two alternating, flashing, images differing in an important detail, interrupted by a blank in between. The change in these movies occurs in a location different from that indicated in Figure S1.*
